# Task-relevant representational spaces in human memory traces

**DOI:** 10.1101/2024.09.10.612205

**Authors:** Rebekka Heinen, Elias M.B. Rau, Nora A. Herweg, Nikolai Axmacher

## Abstract

During encoding, stimuli are embedded into memory traces that allow for their later retrieval. However, we cannot remember every aspect of our experiences. Here, we show that memory traces consist of multidimensional representational spaces whose formats are flexibly strengthened or weakened during encoding and consolidation. In a series of behavioral experiments, participants compared pairs of natural images on either two conceptual or two perceptual dimensions, leading them to incorporate the images into representational “spaces”. We found that distances in task-relevant but not irrelevant spaces affected memory strengths. Interestingly, conceptual encoding did not impair subsequent rejection of similar lures, suggesting that task-irrelevant perceptual information remained in the memory trace. However, targeted memory reactivation following conceptual encoding deteriorated perceptual discrimination, indicating that it weakened the accessibility of perceptual formats. Our results demonstrate that representational formats are flexibly incorporated into memory, and more generally show how the organization of information in cognitive representational spaces shapes behavior.

## Introduction

When you recollect your previous birthday party, you may be able to mentally re-experience numerous details from this event – the taste of the cake, the melodies of the music, and the color of the wrapping paper – and these memories may already fuel imaginations of your next birthday. Episodic memories not only help us remember past experiences but also guide our future behavior. They rely on the reactivation of memory traces – i.e., internal representations of previously perceived events – which can be detected in the human brain via multivariate analysis methods such as representational similarity analysis (RSA) (*1, 2*). These memory traces guide behavioral memory decisions, yet not all information about an episode is stored. Remembering every detail would not only exceed the capacity of the brain, but also contradict the need to generalize past information to future situations. However, it remains unclear which information from an episode is maintained in memory. According to some theoretical frameworks, memory traces are “sparse” and only contain novel or unexpected information, which is complemented at the time of retrieval by pre-existing semantic knowledge (*3, 4*). Alternatively, memory traces may initially contain rich sensory information that degrades slowly over time (*5, 6*). Finally, the amount and kind of information that is stored in memory traces may vary depending on the behavioral goals during an experience (*7*). Here, we sought to test these competing hypotheses on the nature of the memory trace.

During perception, sensory information is hierarchically processed in the brain. For visual content, this hierarchy ranges from low-level perceptual information (e.g., edges and colors) to more complex features (e.g., textures and shapes) to semantic information. Here, we define perceptual and conceptual formats as representations of visual and semantic stimulus features, respectively, thus reflecting how a memory is stored rather than its content. Visual and conceptual features are extracted along the posterior-to-anterior extent of the ventral visual pathway (VVP) and lead to multiple coexisting representational formats of the same stimulus (*5, 8*). A similar processing hierarchy exists in convolutional deep neural networks (cDNN) trained for object recognition, providing a link between representations in different cDNN layers and various processing steps along the VVP (*9–15*). In addition to these sensory formats of increasing complexity, semantic representations can be identified by corpus-based models of deep natural language processing (dNLP) that use text rather than images as input (*16*). Thus, cDNNs and dNLPs provide different quantitative models of cognitive representations across various representational formats (*8, 17–19*). Representational similarities in individual formats correspond to distances in diverse representational “spaces” (with higher distances reflected by lower similarities) that can be employed for cognitive operations such as perception, memory, and reasoning (*20, 21*). While deep neural networks have been shown to model behavioral judgements relying on lower-level representational spaces (*22, 23*) (e.g., defined by animacy or color), it is less clear whether these models can still reliably model representational formats in more complex and abstract spaces (e.g., the approachability of an animal).

While the neural basis and functional role of representational formats has been extensively studied during perception, fewer studies have investigated the formats of memory traces, i.e., of representations in the absence of sensory inputs (*24, 25*). Crucially, certain representational formats may be more relevant for subsequent tasks. These tasks may influence the availability and accessibility of information during later memory retrieval. Using deep neural network models (DNN) models to quantify perceptual and conceptual formats, it was recently shown that conceptual information is preferentially stored in a memory trace and supports later recall, while perceptual information is lost over time (*18, 23, 26*) (for review see (*8*)). This would suggest that memory traces are quickly transformed to only contain conceptual/semantic representational formats.

Prominent theoretical accounts have conceptualized these transformation processes as gist abstraction (*27*) or semantization (*28, 29*). These frameworks propose that transformation condenses detail-rich perceptual contents into more gist-like representations, because conceptual formats may be more generalizable and, therefore, more relevant for future behavior than perceptual formats. Processes of memory transformation and gist extraction have been linked to memory consolidation (*30*), which may start early after encoding (*18, 31*) but is particularly pronounced during sleep (*32*). Consolidation can be experimentally enhanced via targeted memory reactivation, i.e., by presenting auditory or olfactory cues during sleep that have been previously associated with stimuli (*33, 34*). However, few questions remain unanswered. Are perceptual features necessarily lost in favor of conceptual features, or do they remain silently available in the memory trace? In other words, is the representational format indeed subject to transformation (e.g., all sensory information is transformed into conceptual formats), or do multiple formats co-exist with some being gradually strengthened and others gradually weakened? And is this process influenced by task demands?

Indeed, various cognitive theories propose that representational formats depend on behavioral instructions during encoding. This has been suggested both in classical frameworks including levels of processing (*35*) and transfer-appropriate processing (*36*), but also in more recent theories based on reinforcement learning (*37–39*). According to these accounts, task demands during encoding may flexibly determine which representational formats are embedded into a memory trace. Recent work on cognitive maps demonstrates that people mentally “navigate” through abstract, conceptual spaces (*40–44*). Task demands may influence which spaces are recruited and the path taken from one concept to the other may impact subsequent memory.

Further, task demands during retrieval may influence which specific representational format is required. In some cases, it may be necessary to reactivate perceptually detailed information from a memory trace to distinguish previously experienced stimuli from new items that are perceptually similar (pattern separation or differentiation). Other situations may require generalized information about previously experienced categories, independent of the perceptual details of a particular exemplar (pattern completion or generalization). These diverging retrieval demands would benefit from richer memory traces with multiple different representational formats that can be flexibly reactivated depending on situational needs (*23*).

Here, we conducted a series of behavioral studies to address several fundamental questions: How “shallow” or “deep” are representational formats of a memory – i.e., do they contain only a single representational format or multiple formats? How are these formats determined, how may they support different retrieval demands, and (how) do they change after encoding? We investigated the representational formats during encoding, consolidation, and retrieval, and tested (1) whether behavioral goals during encoding affect the representational “space” in which stimuli are embedded, and thus their subsequent memorability; (2) whether metric representational spaces are also employed when participants perform binary judgements during encoding, and whether these abstract spaces can be reconstructed using DNNs; (3) whether different representational formats serve different functional roles during retrieval; and (4) whether the formats that are incorporated into a memory during encoding can be subsequently modified via targeted memory reactivation.

## Results

### Selective embedding of memory traces into task-relevant representational spaces

In studies I and II, participants were presented pairs of natural images of animals and were asked to judge the similarity of these images on either two perceptual dimensions (perceptual blocks), two conceptual dimensions (conceptual blocks), or one perceptual and one conceptual dimension (mixed blocks, only in Study I and removed from subsequent analyses, see Methods; Fig. 1B). During recognition, participants saw all images from the encoding phase together with an equal number of new items (Fig. 1C) and indicated their confidence that a stimulus was old or new on a visual analogue scale (Fig. 1C).

**Fig. 1:**
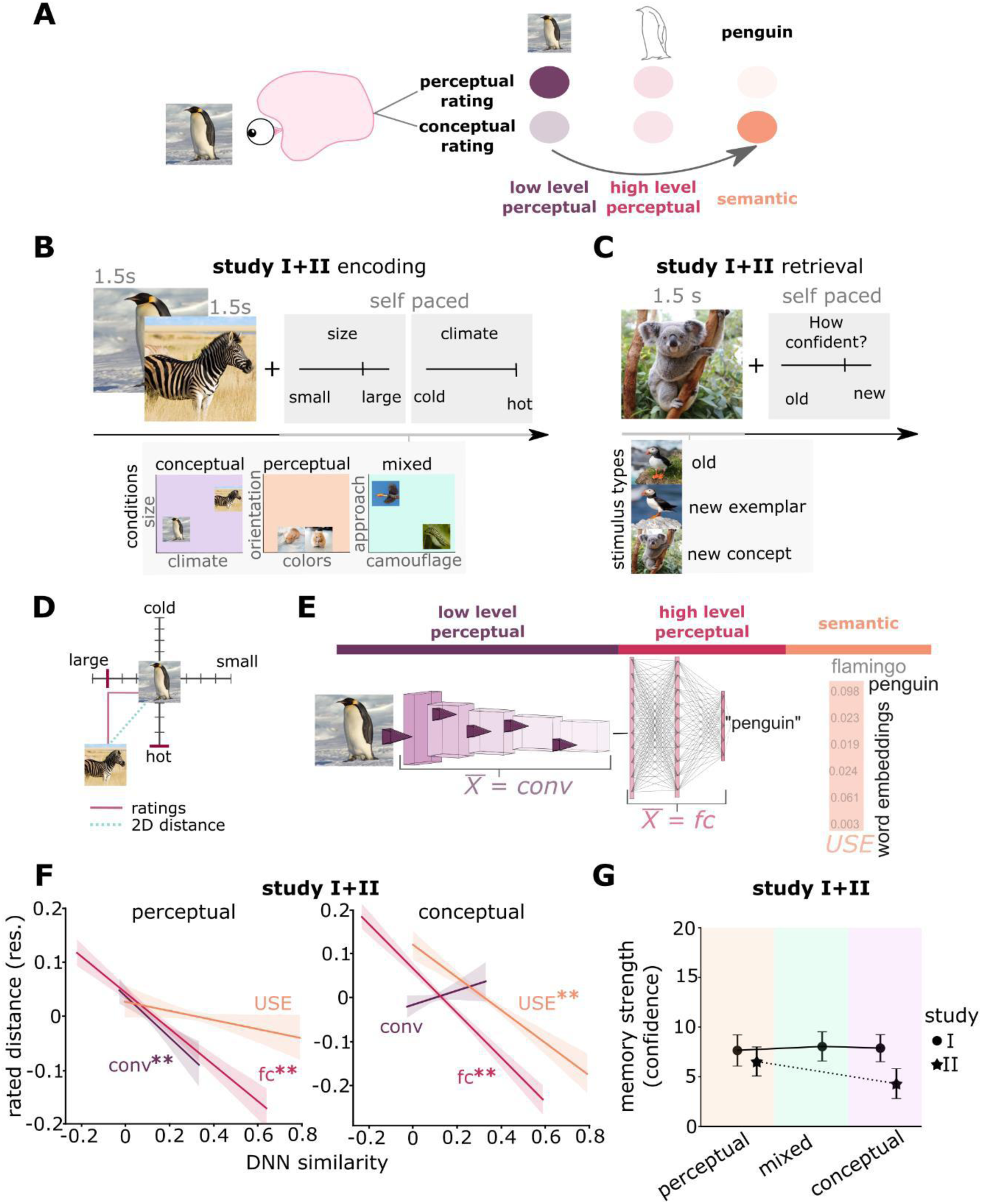
Embedding of task-relevant representational formats into memory traces. (A) Theoretical framework: We hypothesize that memory traces consist of multiple representational formats whose accessibility can be flexibly adapted depending on task-demands. Perceptual tasks may increase the accessibility of perceptual features (left), while conceptual tasks may enhance processes of gist-abstraction and semantization (right). (B) Encoding phase in studies I+II: Participants encoded natural images of animals in either perceptual or conceptual conditions, emphasizing different task-relevant representational formats. In each trial, participants rated the similarity of image pairs on 2 different perceptual and/or conceptual dimensions via visual analogue scales. Study I comprised blocks with 2 perceptual features, blocks with 2 conceptual features, and mixed blocks. Study II contained only perceptual and conceptual blocks, during which 2 different sounds were played for later cueing during sleep (see Figure 4 for more details). (C) Recognition phase in studies I+II: Participants indicated their confidence regarding whether an image was old or new on a visual analogue scale. (D) Representational distances were derived from similarity judgments in each single dimension and from the Euclidean distance of both judgments (2D distance). (E) Representational distances in low-level perceptual formats, higher-order perceptual formats and semantic formats were derived from a convolutional DNN and large language model, respectively. (F) Low-level perceptual and higher-order perceptual similarities predict rated distances during perceptual encoding. Reversely, conceptual ratings can be predicted by higher-order perceptual and semantic similarities. (G) Memory strength did not depend on encoding tasks. Data are visualized after removing participant-wise estimated random effects. 95% confidence intervals, error bars indicate SEM.**, p<0.01

First, we confirmed that our task indeed led to different processing of perceptual and conceptual stimulus formats. To this end, we tested whether similarity judgements in the respective task-relevant perceptual and conceptual spaces corresponded to representational distances in specific DNN layers or models (Fig. 1E). If participants employed perceptual spaces, rated distances should be reflected by distances in convolutional DNN layers that depend on sensory properties, while reliance on a conceptual format should be reflected by distances in a semantic model. Thus, we computed the similarity of each image pair in each network layer of a convolutional DNN (AlexNet(*45*)) and a dNLP (Google’s universal sentence encoder(*46*); USE). We then computed layer averages for the cDNN (see Methods for more details; see Supplementary Fig. 4C for layer-wise correlations) of convolutional (conv) and fully connected (fc) layers. We then used the averaged conv, fc and the USE model similarities as predictors in a linear mixed model on rated distances of Studies I and II including a random participant intercept. Statistical significance was tested using a likelihood ratio test between a full model, including the fixed effect of interest, and a reduced model without the fixed effect of interest (all models included a participant intercept; see Methods for more details; beta weights of full linear models can be found in Supplement Table 1). We found that distances in low-level perceptual space (z = 4.54, χ^2^_(1)_ = 20.67, p < 0.001; likelihood ratio test) and higher-order perceptual space (z = 11.40, χ^2^_(1)_ = 129.51, p < 0.001; likelihood ratio test) but not semantic space (z = 1.87, χ^2^_(1)_ = 3.53, p = 0.06; likelihood ratio test) predicted perceptual ratings (Fig. 1F). Reversely, distances in higher-order perceptual space (z = 15.80, χ^2^_(1)_ = 247.89, p < 0.001; likelihood ratio test) and semantic space (z = 7.30, χ^2^_(1)_ = 53.30, p < 0.001) but not low-level perceptual space (z = 1.76, χ^2^_(1)_ = 3.10, p = 0.07; likelihood ratio test) predicted conceptual ratings (Fig. 1F). Taken together, these results demonstrate that our rating task manipulated participants’ employment of task-relevant spaces. Additionally, the rated distances affected the ease of similarity judgments – i.e., responses to images were given faster when their distances on relevant dimensions were larger (see Supplementary Fig. 2A).

Next, we investigated whether these task-relevant spaces did not only reflect perceptual and conceptual formats during encoding, but whether they were also selectively incorporated into memory traces – i.e., whether the accessibility of individual items for recognition memory judgements was influenced by their task-relevant encoding space. We first analyzed whether the encoding space of an item (perceptual or conceptual) affected the memory strength of that item. Using a linear mixed model of trial-wise memory strength with “encoding task” as a predictor and participant-specific slopes as random effects, we did not find effects for more accessible memory traces following conceptual vs. perceptual judgements (Study I: z = -0.36, χ^2^_(1)_ = 0.42, p = 0.516; Study II: z = 0.15, χ^2^_(1)_ = 0.02, p = 0.874; likelihood ratio test; Fig. 1G). This indicates that conceptual encoding instructions per se did not influence the subsequent accessibility of memory traces, pointing to a possible role of other factors such as the task-relevance of a particular format during encoding.

Next, we analyzed whether memory traces of individual items involved the fine-grained structure of task-relevant representational spaces, by testing whether representational distances in the respective task-relevant format during encoding influenced subsequent recognition memory performance. For each pair of images, we extracted both the task-relevant distances (i.e., the participant-specific ratings of their similarity) and task-irrelevant distances, which were calculated as the average distance of the same pair of images from a least squares solution over all rated images in all other spaces not rated by a given participant (Fig. 2A-B). We then computed a linear mixed model to predict trial-wise memory strengths using the task-relevant distances (2D representational distances) as predictors and participant-specific slopes as random effects. We found that distances in task-relevant but not task-irrelevant formats predicted subsequent recognition memory strength (Fig. 2B). In both Study I+II, higher task-relevant representational distances predicted better recognition memory performance (Study I: z = 4.16, χ^2^_(1)_ = 17.32, p < 0.001; Study II: z = 5.74, χ^2^_(1)_ = 33.02, p < 0.001; likelihood ratio test). In Study II, this effect was significant for both the first (z = -4.65, χ^2^_(1)_ = 21.65, p < 0.001) and second image in a trial (z = -4.46, χ^2^_(1)_ = 19.94, p < 0.001); in Study I, it was significant for the second image (z = -4.27, χ^2^_(1)_ = 18.27, p < 0.001) and by trend for the first image (z = -1.64, χ^2^_(1)_ = 2.71, p = 0.099). Contrastingly, when including the average distance of the same images in task-irrelevant representational spaces as predictors in the linear mixed model, we did not find a significant effect (Study I: z = -0.89, χ^2^_(1)_ = 0.80, p = 0.373; Study II: z = 1.25, χ^2^_(1)_ = 1.58, p = 0.208; Fig. 2b; see Supplementary Fig. 3A-B for separate results in the two encoding spaces). Thus, the irrelevant distances did not predict memory, independent of the processing format. While relevant distances predicted memory confidence for old images, we did not find an effect of relevant distance on correctly rejecting new exemplars as new (Study I: z = -1.51, χ^2^_(1)_ = 2.31, p = 0.128; Study II: z = -0.78, χ^2^_(1)_ = 0.61, p = 0.433).

**Fig. 2:**
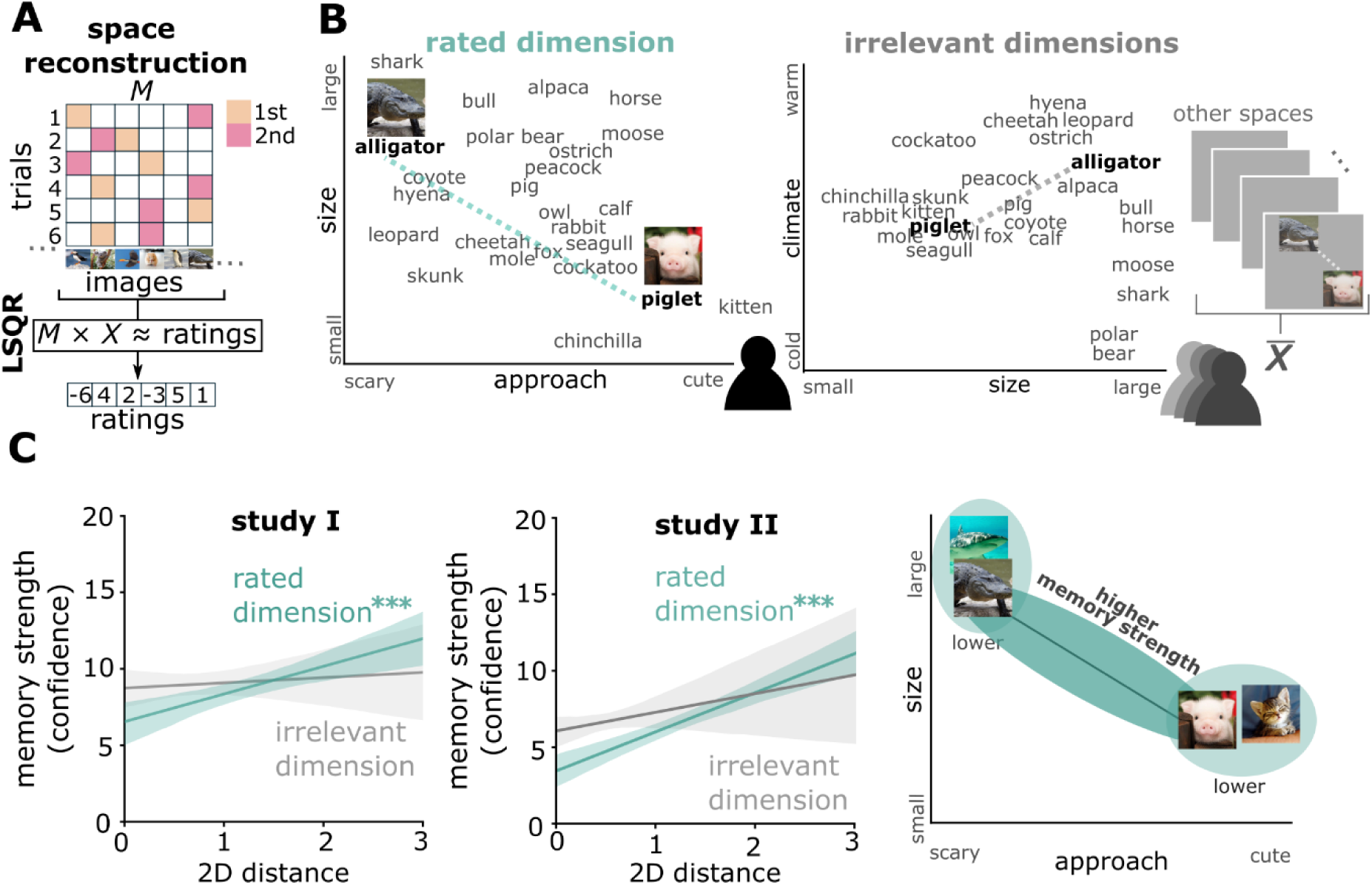
Embedding of task-relevant representational spaces into memory traces. (A) Cognitive space reconstruction. We used the ratings of each participant across all trials and all images to create a parametrization for each rated dimension. A sparse matrix (M) indicating which images were paired was used in a least squares solution (LSQR) given the participant ratings. Thereby each image receives a position value on the given dimension across all participant ratings. We can then combine two solutions (i.e., solution “approachability” and “size”) to create a two-dimensional space. (B) Left: 2D distances in task-relevant encoding spaces were derived from participant-specific similarity judgments (figure shows space in example participant). Right: 2D distances in task-irrelevant spaces were computed as average distance in least square solutions of all other encoding spaces in all other participants (figure shows one corresponding task-irrelevant space across all other participants). (C) Memory strength depends on 2D distances in task-relevant but not task-irrelevant encoding spaces with better memory for higher distances. Left: Study I, Middle: Study 2, Right: schematic depiction. Data are visualized after removing participant-wise estimated random effects. 95% confidence intervals. ***, p<0.001

These results show that task demands not only affect similarity judgements during encoding, but – more interestingly – that they also determine the representational space of an item’s memory trace. Specifically, items that are encoded at higher task-relevant distances are remembered better, possibly reflecting effects of contextual deviance (*47–50*) and/or distinctiveness (*51–53*) (see Discussion).

### Employment of graded and metric representational spaces following binary ratings

In Studies I and II, the representational spaces of item-specific memory traces were defined by explicit distance ratings along graded similarity scales. This might have prompted participants to encode these images into cognitive spaces with metric distances. Next, we tested whether representational distances during encoding also affected memories if participants performed binary decisions in a forced-choice paradigm (Fig. 3A) – indicating the use of cognitive spaces without explicit manipulation. During encoding in Study III, participants first saw a target image and then indicated which of two probe images was more similar to the target image. This decision was again based on either perceptual or conceptual features. We observed high levels of accuracy in selecting the correct choice, with an even higher accuracy for conceptual vs. perceptual ratings (conceptual: M = 0.98, SD = 0.02; perceptual: M = 0.96, SD = 0.03; paired t-test across participants: t_27_ = 4.23, p = 0.0002; the effect size, as measured by Cohen’s d, was d = 0.82).

**Fig. 3.**
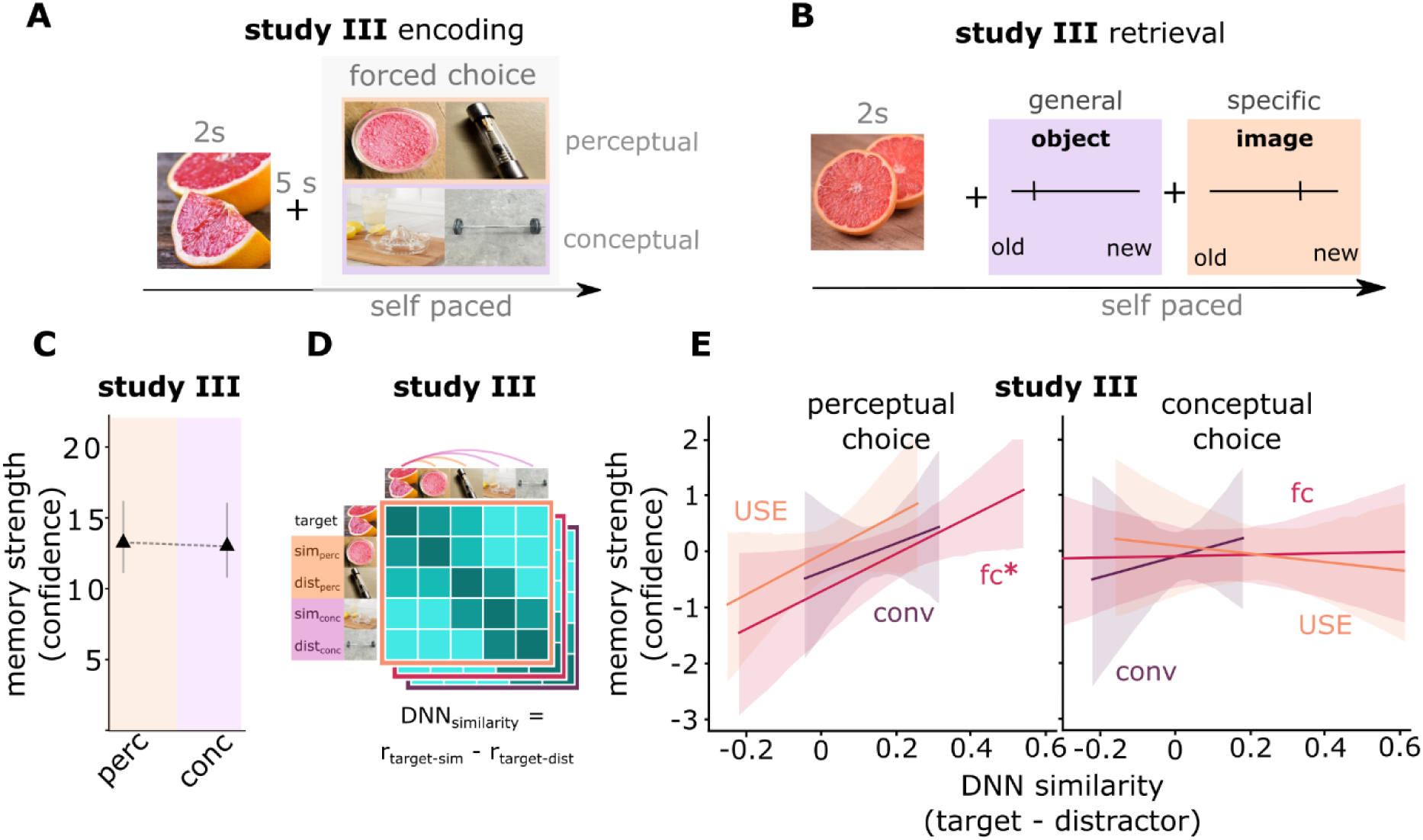
Memory traces with DNN-based representational distances following binary similarity judgments. (A) In Study III, participants performed forced-choice similarity ratings during encoding. Given a target image (blood orange), they chose a conceptually (lemon press) or perceptually (pink power pod) similar image from two options. (B) Participants first performed a general retrieval task (indicating whether a given image matched a previously presented category) and then a specific retrieval task (indicating whether the image matched the previously presented exemplar). (C) Encoding task does not influence memory strength. (D) Graded representational distances in low-level perceptual formats, higher-order perceptual formats, and a semantic format were derived from a convolutional DNN and the universal sentence encoder (USE), respectively. We computed the difference in DNN similarities between target image and similar (correct) choice vs. target and distractor (incorrect) choice, separately for each DNN level. (E) Task-relevant DNN-derived representational distances predict memory strength. Data are visualized after removing participant-wise estimated random effects. 95% confidence intervals, error bars indicate SEM. *, p<0.05.

In order to identify the representational distances that guided participants’ judgements in the absence of graded similarity ratings, we employed DNNs as in Study I and II to extract the similarities between pairs of objects in different representational formats. For each image pair, we thus computed the DNN similarities between the target stimulus and the correct choice image and between the target stimulus and the incorrect lure image (Fig. 3D). The difference between these similarities was considered a proxy for distances in task-relevant representational spaces underlying the decision (see Supplementary Fig. 5 for more details).

Next, we applied linear mixed models to test whether memory traces were embedded into task-relevant representational spaces, i.e., whether task-relevant distances during encoding predicted memory strength. In Study III, we refined the recognition task to separately address different recall demands either based on generalized or pattern-completed information (“Did you see an image from this category during encoding?”), or based on highly differentiated or pattern-separated information (“Did you see exactly this image during encoding?”) (Fig. 3B). Consistent with Studies I+II, we did not find an effect of encoding space on memory strength when using the average confidence of both the specific and general memory test for old images as predictors in a linear mixed model (z = -0.43, χ^2^_(1)_ = 0.19, p = 0.6650; likelihood ratio test; Fig. 3C). Importantly however, for perceptually encoded items, representational distances in higher-order perceptual space (z = 3.91, χ^2^_(1)_ = 5.24, p = 0.022; likelihood ratio test), but not low-level perceptual (z = 2.91, χ^2^_(1)_ = 0.55, p = 0.460; likelihood ratio test) or semantic spaces (z = 3.71, χ^2^_(1)_ = 2.06, p = 0.150; likelihood ratio test; Fig. 3E) predicted memory strength (general and specific memory tests combined; see Supplementary Fig. 6 for separate effects in the general and the specific memory test). For conceptually encoded items, representational distances did not predict subsequent memory strength (all p > 0.597).

Taken together, these results indicate that task-relevant representational distances can affect recognition memory performance, suggesting that memory traces of individual items are embedded in different representational spaces depending on encoding instructions. The linear relationship between task-relevant representational distances during encoding and memory strength (i.e., the accessibility of item-specific information for recognition memory judgements) suggests that memory traces involve coding schemes with metric cognitive maps in specific task-relevant formats (see Discussion).

### Effects of representational formats on different task demands during memory retrieval

We next explored the functional relevance of the different representational spaces for subsequent retrieval demands. Based on previous studies (*54, 55*), we hypothesized that encoding in a conceptual space may be particularly beneficial for retrieval of generalized, perceptually invariant categorical information (*56–58*). Conversely, encoding in perceptual spaces may be relevant when retrieval requires the rejection of similar but not identical exemplars (i.e., pattern separation).

We first analyzed whether the different retrieval instructions in Study III (general/specific) shifted memory decisions (Fig. 4A). Indeed, in the general compared to the specific retrieval task, hit rates were significantly higher (t_27_ = 11.55, p < 0.001; paired t-test; the effect size, as measured by Cohen’s d, was d = 1.50) and correct rejection rates of new concepts were significantly lower (t_27_ = -12.35, p < 0.001; paired t-test; the effect size, as measured by Cohen’s d, was d = 2.59), suggesting a more liberal recognition memory threshold. Note that responses to new exemplars could not be directly compared because they required “old” ratings in the general task but “new” ratings in the specific task.

**Fig. 4.**
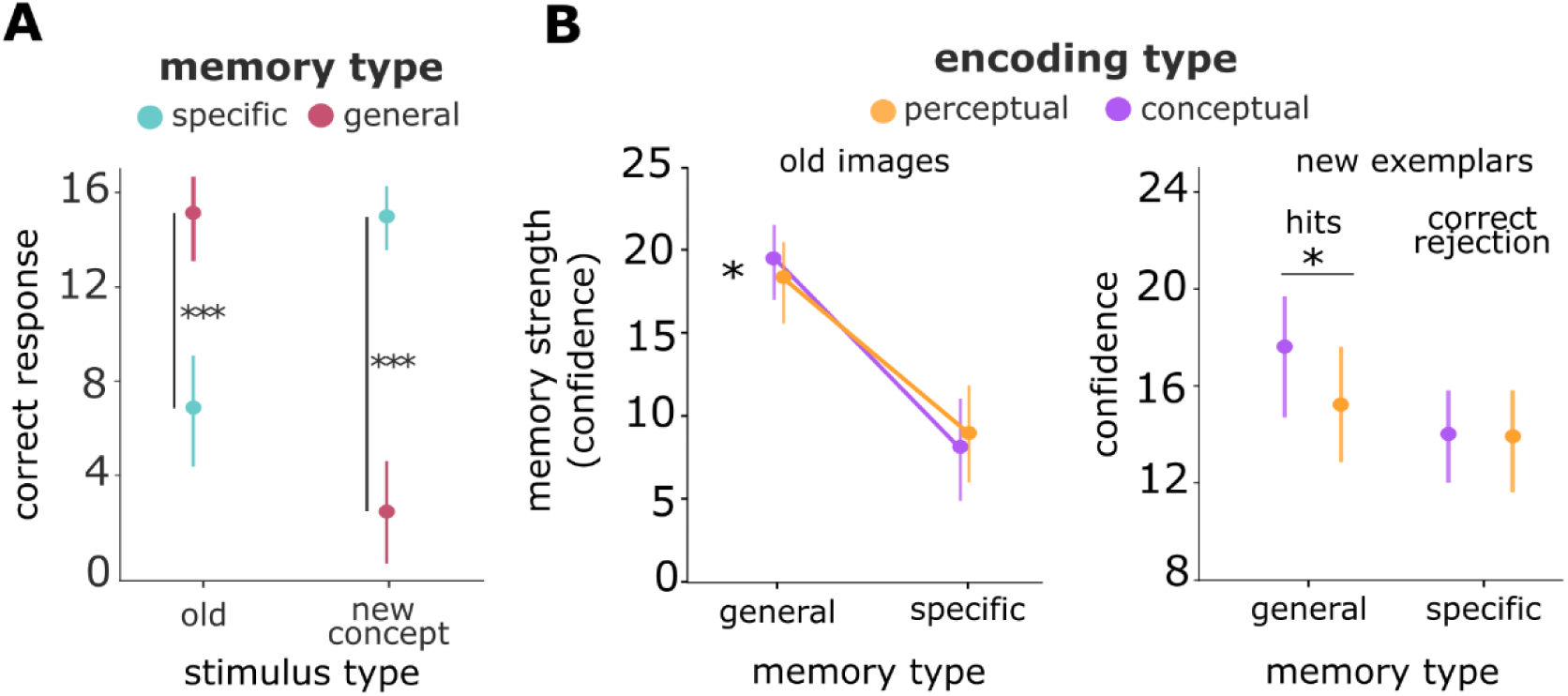
Conceptual encoding improves generalized retrieval and does not impair specific retrieval. (A) Different response criteria depending on retrieval tasks: General retrieval leads to higher hit rates but lower correct rejection rates of new concepts. (B) Left: conceptual encoding improves memory strength during general but not specific retrieval. Right: conceptual encoding facilitates correct recognition that new exemplars belong to previously presented categories during general retrieval and does not influence performance during specific retrieval. Data are visualized after removing participant-wise estimated random effects. Error bars indicate SEM. *, p<0.05, ***, p<0.001

Next, we investigated whether performance in the general and specific memory tasks depended on task-relevant encoding spaces (Figure 4B). We used a linear mixed model to predict memory performance using retrieval task (general/specific) and encoding space (perceptual/conceptual) as predictors with a random participant intercept and including an interaction between retrieval task and encoding space. To test for significant differences, we used a likelihood ratio test between the full model containing the main effect and the interaction term, and the reduced model excluding the interaction. We found an interaction between retrieval task-demands and encoding space (z = 2.69, χ^2^_(2)_ = 7.25, p = 0.007; likelihood ratio test). Conceptual encoding improved memory in the general retrieval task (z = 3.04, χ^2^_(1)_ = 9.27, p = 0.0023; likelihood ratio test) but not the specific retrieval task (z = 1.28, χ^2^_(1)_ = 1.66, p = 0.197; likelihood ratio test; Fig. 4B, left panel). Moreover, we found a significant interaction for new exemplars (z = 2.53, χ^2^_(2)_ = 6.44, p = 0.011; likelihood ratio test). Conceptual encoding benefitted generalized memory decisions, i.e., hit rates towards new exemplars were higher than for perceptually encoded items (z = 4.53, χ^2^_(1)_ = 20.52, p < 0.0001; likelihood ratio test), while performance in the specific retrieval task did not depend on encoding instructions (z = 0.47, χ^2^_(1)_ = 0.22, p = 0.6354; likelihood ratio test; Fig. 4B, right panel). Additional analyses indicated that employment of generalized information (in the general retrieval task) specifically benefited from lower representational distances in higher-order perceptual formats, while rejection of new concepts was impaired by lower distances in this format in both retrieval tasks (Supplementary Fig. 7).

These results indicate that generalized information at test was more readily available for conceptually encoded images for which participants presumably relied more on gist-like representations. In other words, conceptual encoding created memory traces with generalizable representations that facilitated the correct recognition that new exemplars belong to previously presented categories. Thus, encoding stimuli into conceptual spaces improves both the accessibility of their memory traces (higher memory strength of previously presented exemplars) and their employment for generalized memory decisions (better recognition that new exemplars are members of a previously presented category). Interestingly, this does not occur at the expense of less perceptually distinct representations since performance in the specific memory test was not reduced, suggesting that conceptually encoded items are embedded into a “deep” memory trace with multiple different formats (even though task-irrelevant spaces do not seem to involve metric coding schemes; see Discussion).

### Selective strengthening of task-relevant formats during consolidation

So far, our results indicate that memory traces contain encoding spaces in distinct task-relevant representational formats. However, they also reveal an asymmetry, since conceptual encoding spaces benefitted both general and specific retrieval, whereas perceptual encoding spaces did not offer any particular advantage. Since memory consolidation has been suggested to strengthen conceptual representations (*30*), we explored its influence on the representational formats of memory traces. We employed targeted memory reactivation (TMR) and tested whether cueing with a sound that was presented during either perceptual or conceptual encoding selectively enhanced the corresponding representational format (Figure 5A).

**Fig. 5.**
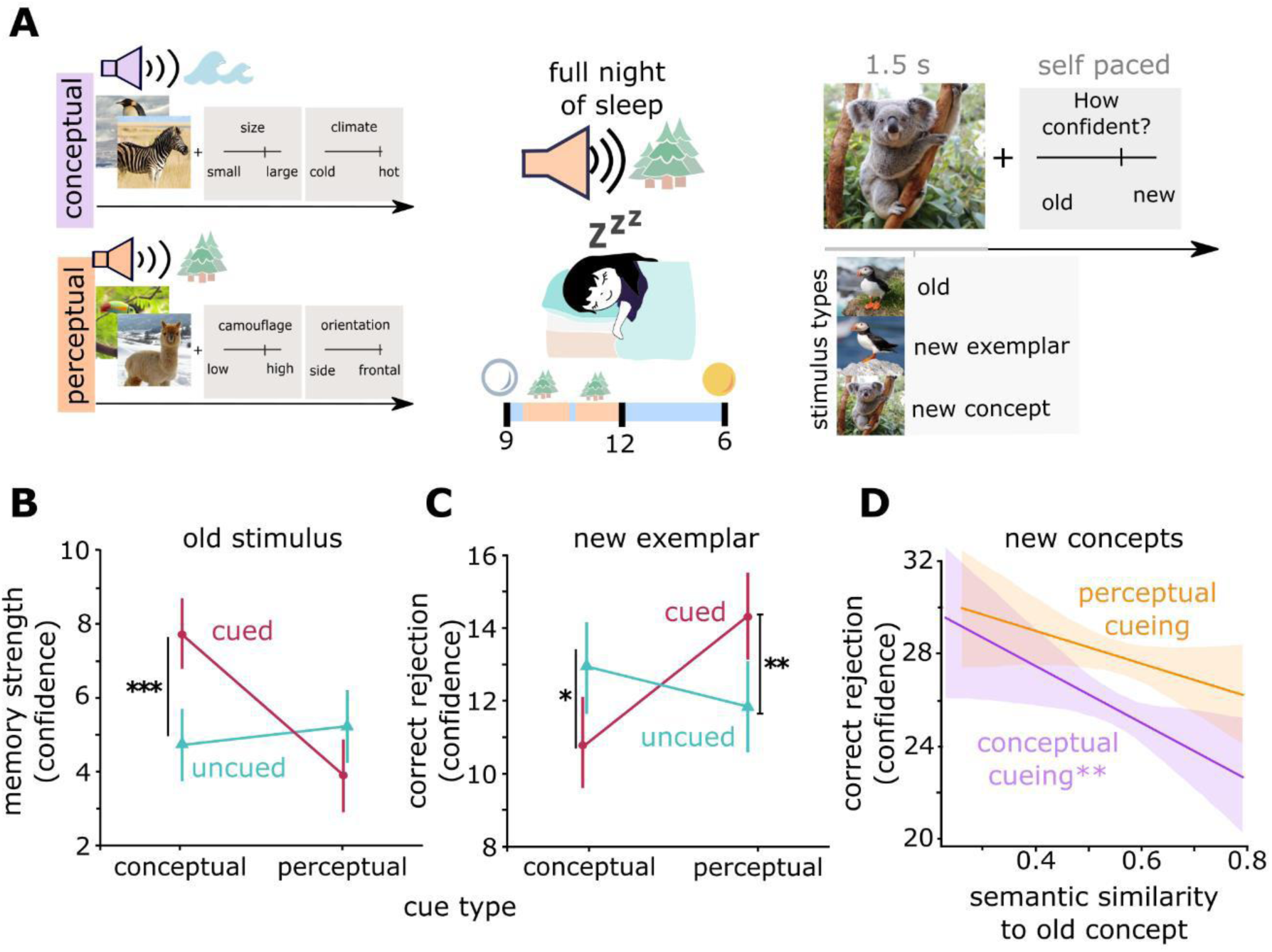
Strengthening of task-relevant and weakening of task-irrelevant representational formats after TMR. (A) TMR design: presentation of condition-specific auditory cues during conceptual and perceptual encoding blocks (e.g., coast sound and forest sound). One of these sounds was played again during the first half of the subsequent night, followed by a graded recognition memory test as in Study I. (B) Cueing of conceptual but not of perceptual formats increases memory strength. (C) Cueing of perceptual formats improves and cueing of conceptual formats impairs correct rejection of new exemplars. (D) Higher semantic similarities of new concepts to previously presented concepts impair correct rejection following conceptual but not perceptual cueing. Data are visualized after removing participant-wise estimated random effects. 95% confidence intervals, error bars indicate SEM. *, p<0.05, **, p<0.01, ***, p<0.001

We first analyzed how TMR affected the memory strength of previously presented items, using a multi-level model with encoding task and cueing as predictors, and participant as random effect (Fig. 5B). We observed a main effect of cueing (z = -2.12, χ^2^_(1)_ = 4.53, p = 0.0333; likelihood ratio test), indicating generally better memory for cued vs. uncued items, but also a significant interaction (z = 4.38, χ^2^_(2)_ = 19.23, p < 0.001; likelihood ratio test), indicating different cueing effects on perceptually vs. conceptually encoded items. Post-hoc tests revealed that cueing only improved the memory strength of conceptually encoded items (z = -4.38, χ^2^_(1)_ = 19.23, p < 0.001; likelihood ratio test) but not of perceptually encoded items, for which even a trend in the opposite direction was observed (z = 1.88, χ^2^_(1)_ = 3.54, p = 0.06; likelihood ratio test).

Since our results thus far demonstrate that memory traces are selectively embedded in task-relevant representational spaces, but also suggest that information in irrelevant formats remains present after encoding, further strengthening one of these formats may come at the expense of other formats. Even though conceptual encoding by itself does not impair specific retrieval of perceptual information (Fig. 5B), one may hypothesize that further strengthening of conceptual formats during consolidation reduces the perceptual specificity of item representations, increasing false alarms to novel exemplars (i.e., lures) of previously encoded categories. We thus implemented a linear mixed model for memory ratings of similar lures with encoding condition and cueing as predictors (Fig. 5C). This model revealed a significant interaction (z = -3.32, χ^2^_(2)_ = = 13.11, p = 0.0003; likelihood ratio test), showing different effects of cueing on correct rejection of similar lures for conceptually vs. perceptually encoded items. Follow-up tests showed that cueing of conceptually encoded items deteriorated the identification of similar lures (z = 2.38, χ^2^_(1)_ = 5.70, p = 0.017; likelihood ratio test), while cueing of perceptually encoded items improved similar lure identification (z = -2.77, χ^2^_(1)_ = 7.66, p = 0.0056; likelihood ratio test). Consistent with our results in Study III, conceptual encoding by itself (i.e., without cueing) did not deteriorate lure detection (z = -0.35, χ^2^_(1)_ = 0.12, p = 0.726; likelihood ratio test). These results indicate that cueing can selectively increase either conceptual representational formats (improving recognition of old items but deteriorating rejection of lures) or perceptual formats (with no effect on later memory but more accurate rejection of lures).

Next, we sought to investigate whether cueing of conceptually encoded items also impairs the ability to identify novel concepts, if they are similar to previously encoded categories. We used the semantic similarities, quantified by the dNLP, of novel concepts to all previously encoded concepts as a predictor in a linear mixed model (Fig. 5D). The results revealed that cueing conceptually encoded items impaired identification of semantically related novel concepts (z = -2.65, χ^2^_(1)_ = 7.01, p = 0.008; likelihood ratio test). This effect was not found after perceptual cueing (z = -1.60, χ^2^_(1)_ = 2.56, p = 0.11; likelihood ratio test). As a control, we confirmed that the influence of low-level perceptual and higher-order perceptual similarities on the identification of novel concepts did not differ between cueing conditions (Supplementary Fig. 8).

These results indicate that representational formats of memory traces are not fully determined by the initial task-relevance during encoding, but that they can be further and selectively strengthened during subsequent consolidation stages. Our findings further show that depending on the task demands to either recognize previously presented items or to reject similar lures, these consolidation processes may exert either beneficial or detrimental effects.

## Discussion

Generative memory theories (*4, 59, 60*) propose that memories are inherently constructive. They suggest that memory traces are rapidly transformed from perceptual into conceptual/semantic representational formats, and that missing information is supplemented from prior knowledge or schemas at the time of retrieval (*3, 61*). Other frameworks propose that memory traces contain richer representations in multiple formats whose relative accessibility can be flexibly determined by various factors during encoding, consolidation, and retrieval (*5, 62*). The aim of our study was to systematically investigate the role of representational formats in memory traces: Are transformations of a memory trace indeed unidirectional, such that perceptual formats are irreversibly lost in favor of semantic information, or do multiple formats coexist? Our study yielded three main results; first, we showed that memory traces consist of distinct representational formats that are determined by task demands during encoding and affect the accessibility of information during subsequent retrieval. The linear relationship between representational distinctiveness and memory strength suggests that stimuli are embedded into metric encoding spaces that are determined by objective item features which correspond to different formats of DNNs. Second, we found asymmetric functional roles of these different encoding spaces for subsequent retrieval demands. While conceptual encoding benefits retrieval during generalized task conditions, it does not impair performance during specific retrieval, suggesting that perceptual information is still maintained in a “deeper”, richer memory trace. Finally, we observed that consolidation selectively strengthens task-dependent representational formats and weakens task-irrelevant formats, leading to “shallower” memory traces.

First, our findings demonstrate the heuristic potential of DNNs to investigate the representational spaces underlying similarity judgements and the selective embedding of these spaces into memory traces. Previous studies have shown that DNNs can be used as a proxy for perceptual and semantic formats during memory tasks (*18, 23, 26, 63, 64*) and behavioral similarity judgements (*64, 65*), and our results indicate that the same models can account for similarity judgements during encoding and their impact on subsequent memory. Importantly, we found that distances from cDNNs and from a natural language model contributed differently. While distances based on perceptual cDNN features predicted perceptual ratings, reaction times, and memory strengths, distances of deeper cDNN layers and of a semantic model predicted behavioral outcomes related to conceptual formats. These results emphasize the usefulness of DNN models as proxies of different representational formats – even in behavioral outcomes – and contribute to the idea of multiple formats in a memory trace.

Contrary to our results, the levels of processing theory suggests a main effect of encoding format on later memory, an effect that we did not find. While unexpected, our results are in line with other studies finding no general effect of processing depth on subsequent memory (*66, 67*). These studies manipulated processing depth using attention orientation tasks (i.e., asking participants to focus on specific stimulus features) (*68*). We used a similar manipulation by drawing participants’ attention to either perceptual or conceptual stimulus features. Interestingly, a recent study showed that image features, measured as reconstruction error of a cDNN, and not the orientation to specific features predicts later memory (*69*). Thus, processing depth may vary based on stimulus features and content rather than processing format (perceptual/conceptual). In addition, the prevalent distance effect in our data may overshadow any main effects of the task.

We observed robust evidence for a role of task-relevant representational spaces. Specifically, the distances of task-relevant but not irrelevant dimensions predicted the memory strength of both perceptually and conceptually encoded items. Thus, task-demands (*7*), but not the mere depth of processing, shapes which features are prone to decay, and which are preserved and even strengthened to support later memory. Interestingly, we found better memory if stimuli were encountered at larger representational distances in their relevant encoding space. How can we explain the higher accessibility of stimuli encoded at higher task-relevant distances? First, one may assume that strengthened memory for more dissimilar items could be caused by higher prediction errors (*70–72*): While the current reference position, e.g., of a shark in the “approachability” space, might lead to the expectation of further dangerous animals, the presentation of a chinchilla would lead to greater prediction errors. However, this explanation cannot account for our finding that not only the second, but also the first image in pairs with larger distances was remembered better. For the same reasons, contextual deviance (as in the von Restorff effect (*50*)) is less likely the main factor driving this result (*73, 74*). Alternatively, higher dissimilarities may promote processes of pattern separation (*75*), which may further facilitate the disambiguation of similar images during retrieval. However, we did not find that higher task-relevant encoding distances facilitated the correct rejection of similar lures. Previous studies showed that the hippocampus encodes representational distances in task-relevant cognitive maps (*41, 76*). Here, we extend these findings by showing that these distances are actually relevant for subsequent memory, and thus contribute to theories bridging spatial navigation and episodic memory (*20, 77*). Specifically, we found that representational distances were linearly related to the accessibility of items for later memory, even if items were not originally encoded using graded distance ratings (Study III). This suggests that participants employed cognitive maps with interval-scaled (metric) rather than rank-based (ordinal) coding schemes (*78*), in line with the original concept of cognitive maps to promote inference during physical navigation (*79*).

Notably, while all three studies presented here provide converging evidence for a selective strengthening of task-relevant but not task-irrelevant representational formats, we also consistently found that conceptual encoding by itself did not come at the expense of reduced identification of similar lures. This suggests that the benefit of conceptual encoding may not (only) lie in deeper encoding and faster integration with pre-existing semantic knowledge, but (also) in the formation of “deeper” and more versatile memory traces that can be flexibly employed for both specific and generalized retrieval demands. Even though the accessibility of perceptual information in conceptually encoded memory traces does not depend on (in this case, task-irrelevant) perceptual distances, and is thus not organized in a “metric” cognitive map (see above), it remains available even following a night of sleep (Study II). Future studies will need to investigate whether more extended consolidation processes across longer time periods promote the formation of “shallower” traces in purely conceptual/semantic formats, leading to a more pronounced decline in the availability of perceptual information.

Interestingly, and to our knowledge for the first time, we observed detrimental effects of conceptual as compared to perceptual encoding on lure identification following TMR, suggesting that enhanced consolidation processes induce a “pruning” of memory traces that removes information in task-relevant representational formats. Previous studies have demonstrated that cueing during sleep is associated with reactivation of the associated memories (*80–84*). After cueing conceptually encoded items, participants relied more on gist-like representations, causing a decreased performance in the rejection of both visually similar lures and semantically similar concepts. These findings are in line with results that reactivation during sleep facilitates generalization processes of features that are shared across several encoded contents (*85–87*), which in turn increases false alarm rates for similar lures (*88*). However, while this may be interpreted as evidence for a unidirectional transformation of memory traces from perceptual into conceptual formats during consolidation, we also found that cueing of perceptually encoded items improved the correct rejection of visually similar lures. This suggests that TMR – and possibly, memory consolidation and sleep-related reactivation more generally – selectively enhances task-relevant (*33, 89*) and reduces task-irrelevant representational formats, rather than generally transforming perceptual into conceptual formats. Recent work indeed suggests reactivation-induced forgetting during sleep: While target memories were enhanced, this enhancement was accompanied by forgetting of irrelevant competing memories (*90, 91*). Thus, while multiple representational formats coexist in memory traces after encoding, these formats can be selectively strengthened and weakened by consolidation processes (*5*).

Our results are solely based on behavioral measures. While this does not allow us to directly demonstrate representational formats of memory traces on a neural level, we understand behavior as a measurable expression of the underlying memory trace (*1*). Additionally, our study emphasizes the utility of behavioral paradigms during (arguably) ecologically valid tasks for inferring complex properties of human cognitive representations (*92*). Nevertheless, we propose that follow-up studies should aim to investigate these representations more directly using neuroimaging and/or electrophysiological recordings. Specifically, previous studies have emphasized a role of the hippocampus for learning (*93*) and for representing multidimensional abstract spaces (*20, 41, 94*). Thus, it would be interesting to investigate whether hippocampal circuits can be flexibly adapted to represent the respective task-relevant formats due to their high degrees of mixed selectivity (*95*), or whether different formats are represented in different hippocampal regions (e.g., along its long-axis) (*96–98*). Moreover, while previous studies investigated representations in newly formed abstract spaces (*40, 99, 100*), participants in our study performed similarity and memory judgements in pre-existing spaces of semantic and perceptual relationships, which may involve interactions of hippocampal distance representations with distinct regions along the VVP. Finally, neuroimaging studies are ideally suited to investigate “dormant” memory traces during states without direct behavioral readouts, such as awake/resting periods, sleep, or while concurrent cognitive tasks are performed. Thus, these studies can explore the mechanisms underlying the “pruning” of memory traces during consolidation.

Taken together, we propose that both perceptual and conceptual representational formats coexist with various factors such as attention, prior knowledge, and task-demands determining the accessibility of a specific format thereby shaping the apparent transformation of memories over time and contexts.

## Materials and Methods

### Sample

We tested healthy adult students from Ruhr University Bochum (Germany). Participants gave written informed consent and received study credits as compensation. All three studies were approved by the institutional review board of the Faculty of Psychology, Ruhr University Bochum (Study I+III: #730, Study II: #554), and were conducted in accordance to the declaration of Helsinki. In Study I, we tested 47 participants (age M = 22.9, SD = 4.0, 38/9 female/male). In Study II, the sample comprised 55 participants (age M = 23.05, SD = 3.57, 44/11 female/male) after excluding one participant who woke up during the night and recognized the sound played. In Study III, we tested 28 participants (age M = 23.17, SD = 3.22, 16/12 female/male) after excluding three participants due to technical issues. Analysis of DNN contributions to prediction of encoding distances was done on the combined data from Studies I and II (n = 102).

### Behavioral Tasks

Data for all three studies were collected online. We implemented the tasks for Studies I and II using jsPsych (*101*) and Cognition (https://www.cognition.run/). In Study III, we implemented the task using Psychopy3 (*102*) and Pavlovia (https://pavlovia.org/).

We chose images from the THINGS database (*103*), a large database of natural images from different categories (e.g., animals, objects, and food), specifically built to make DNN training sets more naturalistic and applicable to neuroscience. Items in this database represent an ecologically valid reflection of human-relevant everyday objects. For Studies I and II, we used animal images only to avoid effects of animate/inanimate differences during later similarity analyses. We excluded images containing multiple animals or containing humans. Since the encoding phase of Study III consisted of predefined forced choice trials, the conceptual and perceptual triplets (target and two choice options) showed not only animals, but equal amounts of objects from other categories contained in the THINGS database, namely food, tool, container, vehicle, and a compilation of individual objects from miscellaneous categories (e.g., musical instruments, electronic devices, and toys). For each participant, the amount of target objects from each category was kept equal within encoding blocks and across perceptual and conceptual conditions. During recognition, new concepts were selected from the same pool of categories and in equal amounts.

Studies I and II consisted of two parts, an incidental encoding task (Fig. 1B) and a recognition task on the following day (Fig. 1C). During encoding, participants rated conceptual or perceptual similarities of a series of image pairs drawn from different categories. In each trial, we consecutively presented two images for 1.5 seconds each, followed by two self-paced ratings which required either conceptual or perceptual similarity judgements. Participants rated similarities of the last presented image in relation to the first image on a visual rating scale. Conceptual judgements included size (“Is the animal smaller or larger?”), climate (“Does the animal live in a colder or a hotter climate?”), and approach (“Is the animal more approachable or less?”). Perceptual judgements included orientation (“Is the animal more facing towards the viewer or more away from the viewer?”), colors (“Does the image contain warmer or colder colors?”) and camouflage (“Is the animal concealed more or less?”). Since we wanted to examine if perceptual or conceptual formats, or the combination of both, would strengthen subsequent memory, we divided the six rating dimensions into three different block types. Each type consisted of two dimensions from either the same or different conditions (perceptual, conceptual, mixed). We presented 336 images of 84 animals in total, with each animal being shown in one of the three block types only (e.g., the four different images of giraffes were all shown in perceptual blocks). Block types of each animal and image pairing were randomly shuffled between participants.

In Study II, we investigated the role of consolidation on the transformation of representational formats. We used the same design as in Study I but paired conceptual and perceptual ratings with condition-specific sounds (Fig. 5A). During each encoding block, we continuously played one of two distinct nature context audios (*104*). One audio was composed of ocean waves and sounds of the sea. The other audio consisted of forest sounds, like the brushing of leaves in the wind, frogs croaking, and crickets chirping, excluding bird sounds to avoid interference with our animal classes. One of the two sounds was played again during post-encoding sleep to reactivate memories associated with the sound (*105, 106*). Each audio file was randomly assigned to be played either during conceptual or perceptual encoding blocks. We decided to exclude mixed rating blocks, as there would be no clear conclusion regarding how consolidation of both rating dimensions should impact subsequent memory performance, while simultaneously, this would have increased the number of trials to cue during sleep. In the night following the encoding session, participants were asked to play a custom-made audio file using their smartphone, starting the file when they would start trying to fall asleep for the night. In standardized studies applying targeted memory reactivation, EEG data would be monitored online to specifically play sounds during slow wave sleep (*33, 107*). Due to practical reasons and to ensure contact-free participation in our study during the COVID-19 pandemic, we decided to play an audio file timed in accordance with the expected higher amount of slow wave sleep during the first half of the night (*108–110*). Audio files were composed to play either the perceptual or conceptual sound during the first half of sleep with long periods of slow wave sleep, which are highly relevant for consolidation (*107, 109*). The audio file started with a masking white noise to habituate participants to the volume, in order to avoid waking them up due to the sudden onset of the actual cue playing during sleep. Sixty minutes after audio onset, the cue was played for 30 minutes, followed by 60 minutes of masking noise. The cue was then played again for 30 minutes. If participants had been awake for longer periods or had to use the bathroom until 2 a.m., they were instructed to restart the audio, though no participant reported waking up prior to this time. The next morning, participants were asked whether they heard any specific sounds during sleep, woke up, or stayed awake for longer time periods. If participants recognized the sounds being played during sleep or if they did not have a full night of sleep between encoding and recognition, we excluded these participants from later analysis (n = 1). We also excluded trials for Studies I and II with reaction times greater than 150 seconds.

The day following encoding, participants performed a recognition memory task which was identical for Studies I + II (Fig. 1C). We presented the same images again (“old”), mixed with a total of 336 new images. Novel images were either unknown images of previously presented animal species (“new exemplars”) or images of animal species that had not been presented before (“new concepts”). Each image was presented for 1.5 seconds. Afterwards, participants indicated their confidence of having seen the image before on a continuous rating scale from old to new. All 336 old images, 168 new exemplars, and 168 new concepts were shown in a random order.

To examine the role of binary rather than gradual similarity ratings and of different retrieval demands, we implemented different encoding and retrieval tasks in Study III (Fig. 3A-B). Instead of subjectively rating the similarity of image pairs during encoding, participants performed forced-choice similarity judgements of triplets (three serially presented images). Each triplet began with a 2s presentation of the target image, followed by a 5s maintenance period during which a fixation cross remained on screen. After maintenance, two horizontally arranged images were presented on either side of the fixation cross (left and right choice option). Participants were instructed to select the image that was more similar to the initially presented target on either perceptual or conceptual dimensions. Similar to Studies I and II, participants performed blocks of the encoding task in which either perceptual (image) or conceptual (object) features were relevant for performing the task.

In order to ensure reliable and valid responses across participants, we carefully constructed trials for each target image where either a perceptually or conceptually similar option was shown together with an unrelated distractor. Stimuli were selected based on similarity predictions from the cDNN. For each of the 120 target images, we manually selected images to form a conceptual and perceptual triplet by identifying images from the THINGS database which predominantly shared feature similarities with the target on either early (convolutional) DNN layers (perceptual, e.g., similar colors, shapes: blood orange - pink powder) or later (fully-connected) DNN layers (conceptual, e.g.: similar object category: blood orange - lemon press). Accordingly, for each triplet, we selected distractor images that did not share the respective features with the target image. To ensure a meaningful difference in similarity of target and choice options, we quantified the difficulty of each triplet by computing the difference in the similarity between the target and the matching stimulus (r_sim_) and the similarity between the target and the distractor (r_dist_). For the distribution of scores on each visual and semantic DNN layer, see Supplementary Figure 4. We restricted our analysis to triplets with >75% accuracy across participants, which led to exclusion of n = 5 perceptual triplets.

After successfully performing the forced choice task, participants received monetary reward feedback of two cents (low) and 10 cents (high). Reward conditions did not elicit behavioral differences across conditions in neither encoding or recognition, nor their congruency in encoding-recognition, and are thus not further discussed. For all analyses of Study III encoding, N = 3,161 trials were considered after excluding trials with incorrect forced choice response (n = 86) and trials with reaction times exceeding four standard deviations above the participants’ average (n = 44).

Similar to Studies I+II, participants engaged in a recognition memory test on the following day (Fig. 3B). During this test, 120 old images, 120 new exemplars, and 120 new concepts were presented for two seconds each. In each trial, participants consecutively indicated their confidence of having seen the object (general retrieval) and having seen the specific image (specific retrieval) using two separate scales. Participants indicated general memory before specific memory, similar to the procedure in (*23*). Note that while for old images (old concept, old image) and new concepts (new concept, new image), correct responses in the two retrieval tests were on identical sides of the slider scale, this differed for new exemplars, which required “old” responses in the general retrieval task but “new” responses in the specific retrieval task.

As for the encoding task, we excluded trials with reaction times exceeding four standard deviations above the participants’ average reaction time (n = 120 perceptual trials, n = 120 conceptual trials), resulting in a total trial number of N = 9,494 recognition trials across all participants.

### Linear mixed models

We analyzed all data from the three studies using linear mixed models of the statsmodels package (*111*) in Python 3.8. To determine the influence of different factors on behavioral performance, we set up a full model containing all fixed effects, and participant as a random effect. We then used a likelihood test to determine whether the exclusion of the factor of interest significantly reduced the explained variance compared with the full model. For main effects with one predictor, the reduced model included a constant as fixed effect (1). For main effects with two or more predictors, the reduced model contained all other main effects except the effect of interest. For interaction effects, the full model contained all main and interaction effects. However, main effects from interaction models were not interpreted. For each fixed effect, we report z-statistics from the initial estimation of the model and the respective likelihood derived from model comparison (χ2) and its significance level (p). Beta weights resulting from the full models are shown in Supplementary Table 1 (for the main results) and Supplementary Table 2 (for analyses from the Supplement). For post-hoc estimation of condition differences, we ran separate models on data split by the factor of interest, and then compared the likelihood of the reduced model with only a random intercept. For visualization purposes of individual fixed effects, we plotted the residuals of mixed models missing the fixed effect of interest.

Finally, individual models were used to determine the behavioral performance in conceptual vs. perceptual encoding tasks as well as memory performance of old images, new exemplars, and new concepts during recognition.

### Task-relevant versus task-irrelevant distances

To compare the effect of relevant and irrelevant distances in Studies I+II, we first constructed individual one-dimensional spaces for each dimension by computing a least squares solution on a sparse matrix of all image ratings on the respective dimension over all participants (Figure 2A). For this, we first created a design matrix M with dimensions of the number of ratings by the number of images containing zeros. Next, M was filled for each rating with 1 or -1 indicating if an image was first (1) or second (-1) in the rating trail. The design matrix M was then converted into a compressed sparse column matrix. A least-squares solution was then used to solve the linear system of equations on our design matrix M given the ratings done by participants. Two least squares solutions can then be used to compute the distance of each image pair in the respective 2D space. We then use these spaces to extract the distances of image pairs on dimensions that the participant did not rate the image pair on, thus creating an “irrelevant” distance as an average over all spaces. The irrelevant distance in this case thus corresponds to a distance that was not relevant for the participant during the task. In contrast to this, we define the relevant distance as the similarity judgment of each image pair that the participant performed during the task. For example, when a participant rated the similarity of a panda and a giraffe on the dimensions of climate and size (rated distance), we used the average distance from the least square solutions of all other spaces (climate + approachability; size + approachability; color + camouflage; orientation + camouflage; color + orientation) as the irrelevant distance.

### DNN similarities

We used a pre-trained convolutional deep neural network, “AlexNet” (*45*), as implemented in the Caffe framework (*112*) in Python 2.7, and abstract semantic representations from the Google Universal Sentence Encoder (*46*) using tensorflow (*113*) in Python 3.8. (Fig. 1E). To analyze cDNN similarities, we first generated features for each layer of the AlexNet for all images. In each layer we averaged across the spatial dimension, retaining just one value per feature. We then correlated the features of each image with all other images using spearman’s *rho* correlation, resulting in eight image-by-image correlation matrices, one for each layer. These correlation maps represent the similarity of each image to all other images across the layers of the cDNN, starting with similarities based on early visual information in convolutional layers and concluding with higher-order visual information of fully connected layers (Fig. 1E). We averaged the correlation maps of convolutional and fully connected layers to simplify later interpretation of differences concerning behavioral performance and reducing the number of statistical tests to correct for. Thus, we obtained two correlation matrices from AlexNet, which served as our markers of low-level perceptual representations (convolutional layers) and higher-order perceptual representations (fully connected layers), while the USE model contains abstract semantic feature formats. We computed word embedding vectors for all concept labels retrieved from the THINGS database and correlated each concept embedding vector with all others to generate a concept-by-concept correlation matrix.

Pairwise image similarities from perceptual and semantic DNNs were used to quantify the degree of overlap between pairwise images, given specific features (Supplementary Fig. 4). For Studies I+II, we capitalized on the behavioral ratings. Participants were instructed to rate the relation of both images on two dimensions, where the second image was rated relative to the first image. We expected the effect to be dependent on the rated dimension (conceptual vs. perceptual) with higher contributions of early (convolution) cDNN layers to perceptual ratings, and of late (fully connected) cDNN layers and USE to conceptual ratings. This analysis was done on the layer-wise correlation maps to investigate the effect in more detail across the DNN gradient. For each layer, we performed a one-sided paired t-test (scipy package (*114*)) on fisher-z transformed average correlations between participant ratings of perceptual and of conceptual dimensions (conceptual, perceptual, and mixed for Study I; conceptual and perceptual for Study II) and the similarities of the respective cDNN layer and USE, resulting in nine paired t-tests. We then corrected for multiple comparisons using FDR-correction. We computed the difference between perceptual and conceptual similarities in each layer to plot layer-wise relations to similarity judgements. Next, we used layer-averaged, and fisher-Z transformed similarity matrices as predictors in mixed models for rated distance during encoding, and for memory strength during recognition.

To determine the degree of format-specific processing during memory formation, we constructed trial-specific measures indicating the distinctiveness of similarity judgements of task-relevant feature formats. Each image was considered a point in 2D space, of which we computed its trial-specific Euclidean distance. The first image was the point of reference with coordinates [0,0], while the coordinates of the second image were the given ratings on each scale [rating1, rating2]. For Studies I and II, we computed the similarity of a new exemplar or new concept to all images previously shown during encoding on conv and fc layers for perceptual similarity and the similarity of a new concept to all previous concepts and extracted the highest similarity for later analyses. For Study III, we computed the similarity of a new exemplar to its matched images during encoding. For new concepts, we extracted the highest similarity of a new concept to all old concepts from encoding.

In Study III, we extracted our measure of distinctiveness from the pairwise similarities across DNN-derived layer predictions for all three images of a triple (target, similar choice, distractor choice) (see Supplementary Fig. 5). We took the similarity of the target image with the similar choice (r_sim_) and the distractor choice (r_dist_) and used their difference (d = r_sim_ - r_dist_) as quantification for triplet (dis-) similarity. Both measures of trial-distinctiveness quantify the degree of difficulty during a given trial (e.g., small distances of animals indicate similar features), making the magnitude and direction of the given rating more difficult. Likewise, low triplet-distinctiveness indicates that both choice options (similar and distractor) are approximately equally similar to the target image, making the forced choice task more difficult. This was done for all images of a triplet in Study III and all pairs of old images and their corresponding new exemplars. This was also done to estimate the distance of new concepts to the bulk of all previously seen concepts by fetching the highest similarity to all old concepts (i.e., smallest distance).

## Data and material availability

All data needed to evaluate the conclusions in the paper are present in the paper and/or the Supplementary Materials. Data and analysis code to reproduce the main results presented in the manuscript are publicly available at (https://osf.io/qn2gw/?view_only=2ca202d43ad8412e86282e020d78068f).

## Supplementary Materials

**Supplementary Fig. 1.**
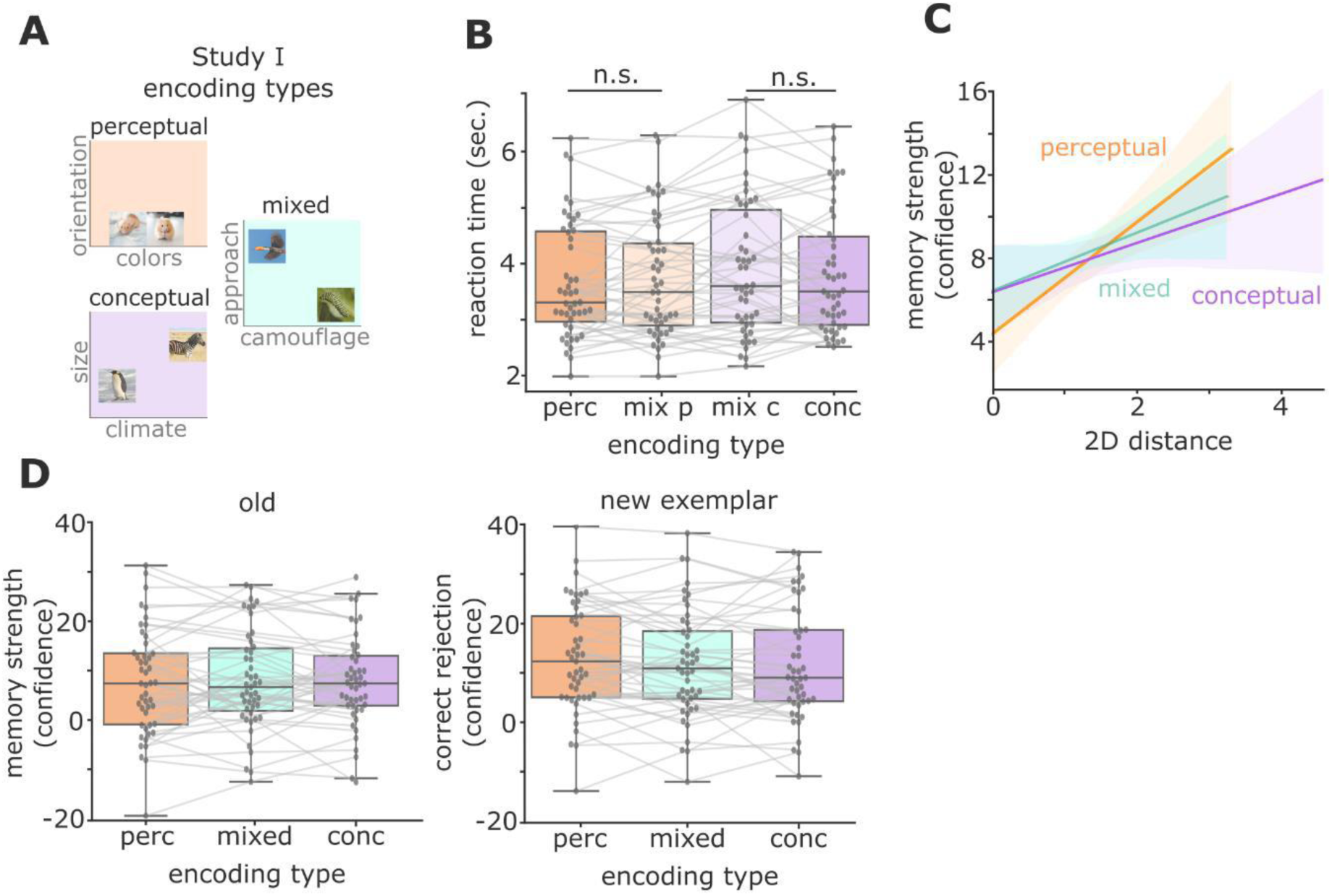
Mixed blocks reflect perceptual and conceptual formats on single dimension levels: (A) In Study I, we presented perceptual blocks (two perceptual dimensions), conceptual blocks (two conceptual dimensions), and mixed blocks (one perceptual and one conceptual dimension). We investigated whether ratings in mixed blocks would differ from ratings in congruent (i.e., purely perceptual, and purely conceptual) blocks. (B) We used linear mixed model to predict reaction times using encoding type (perceptual/conceptual/mixed) as a predictor, including a random participant intercept. Reaction times during perceptual ratings in mixed blocks (z = -1.65, χ^2^_(1)_ = 2.72, p = 0.098; likelihood ratio test) and during conceptual ratings in mixed blocks (z = 1.38, χ^2^_(1)_ = 1.91, p = 0.167; likelihood ratio test) did not differ from their congruent counterpart. (C) We then assessed whether the influence of rated distances on memory strength differs between encoding spaces. Using a linear mixed model including the interaction of encoding space (conceptual, perceptual, mixed) and 2D distance as predictors, we found no interaction of 2D distances and encoding space (perceptual vs conceptual: z = 1.05, mixed vs conceptual: z = 1.04, mixed vs perceptual: z = 1.38, χ^2^_(2)_ = 2.89, p = 0.235; likelihood ratio test). (D) Memory strength (left; perceptual versus conceptual: z = 0.49, mixed vs conceptual: z = -0.11, mixed vs perceptual: z = 0.64, χ^2^_(1)_ = 0.42, p = 0.810; likelihood ratio test) and correct rejection of new exemplars (right; conceptual vs perceptual: z = -1.74, mixed vs perceptual: z = -1.61, mixed vs conceptual: z = -0.10, χ^2^_(1)_ = 3.78, p = 0.151; likelihood ratio test) did not differ between blocks. Based on these results, we decided to concatenate perceptual dimension ratings from mixed and perceptual blocks and conceptual ratings from mixed and conceptual blocks for further analysis of DNN-based similarities in Study I. Data are visualized after removing participant-wise estimated random effects. 95% confidence intervals, center line indicates median, box limits indicate upper and lower quartiles, whiskers indicate 1.5 interquartile range, individual points represent participant averages. n.s., not significant

**Supplementary Fig. 2.**
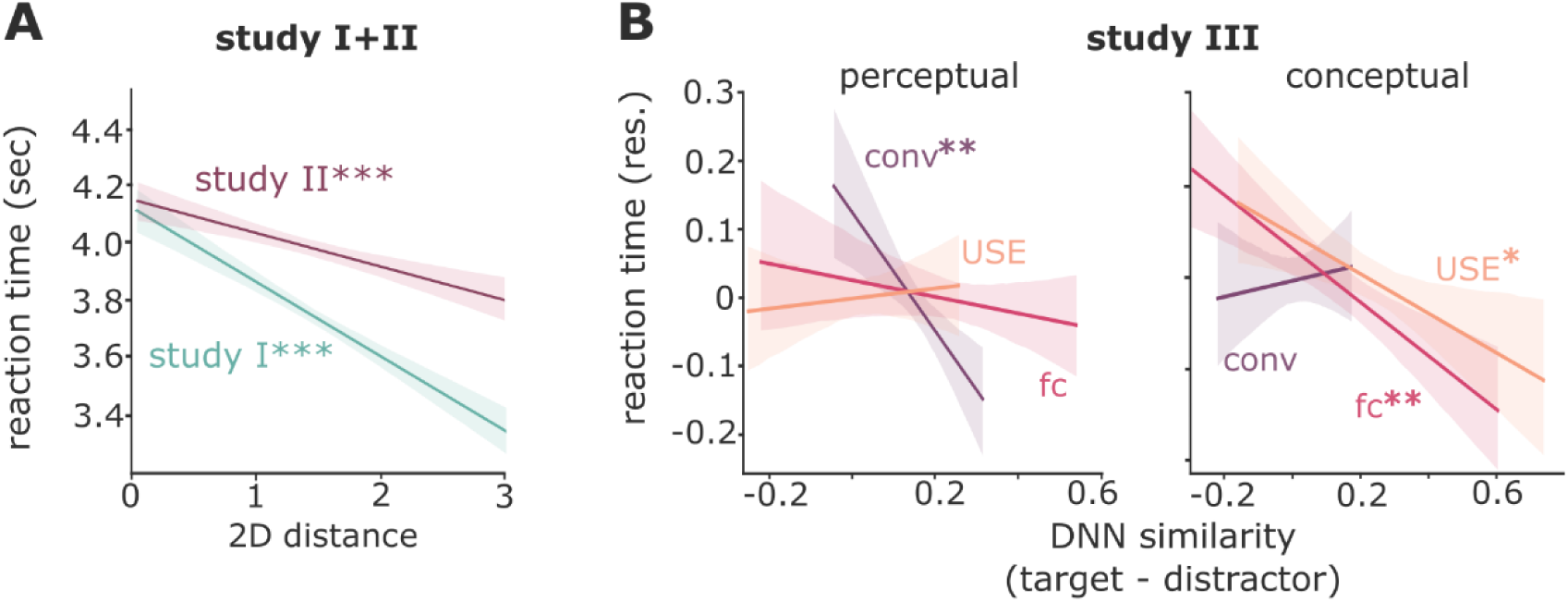
Reaction times are influenced by rated distances: (A) We first tested whether representational distances affected the ease of similarity judgments – i.e., whether responses to images were given faster when their distances on relevant dimensions were larger. We used a linear mixed model to test the effect of task-relevant distance on reaction times including a random participant intercept. Significance testing was done using a likelihood ratio test between a full model, including the fixed effect of interest, and a reduced model without the fixed effect of interest (participant intercept only). Unsurprisingly, we found that larger 2D distances of task-relevant dimensions predicted faster response times (Study I: z = -11.28, χ^2^_(1)_ = 126.69, p < 0.001; Study II: z = -6.67, χ^2^_(1)_ = 44.49, p < 0.001). (B) Similar to Studies I and II, we found higher distances to predict faster response times. Response times during perceptual choices were predicted in a linear mixed model with a random participant intercept using distances in low-level perceptual space (z = -3.90, χ^2^_(1)_ = 15.20, p = 0.001; likelihood ratio test) but not higher-order perceptual space (z = -1.29, χ^2^_(1)_ = 1.67, p = 0.196) or semantic space (z = 0.46, χ^2^_(1)_= 0.22, p = 0.639; likelihood ratio test). Contrastingly, response times during conceptual choices were predicted by distances in both higher-order perceptual (z = -4.91, χ^2^_(1)_ = 24.00, p < 0.001; likelihood ratio test) and semantic spaces (z = -2.68, χ^2^_(1)_ = 7.21, p = 0.007; likelihood ratio test), but not low-level perceptual space (z = 0.49, χ^2^_(1)_ = 0.24, p = 0.621; likelihood ratio test).

**Supplementary Fig. 3.**
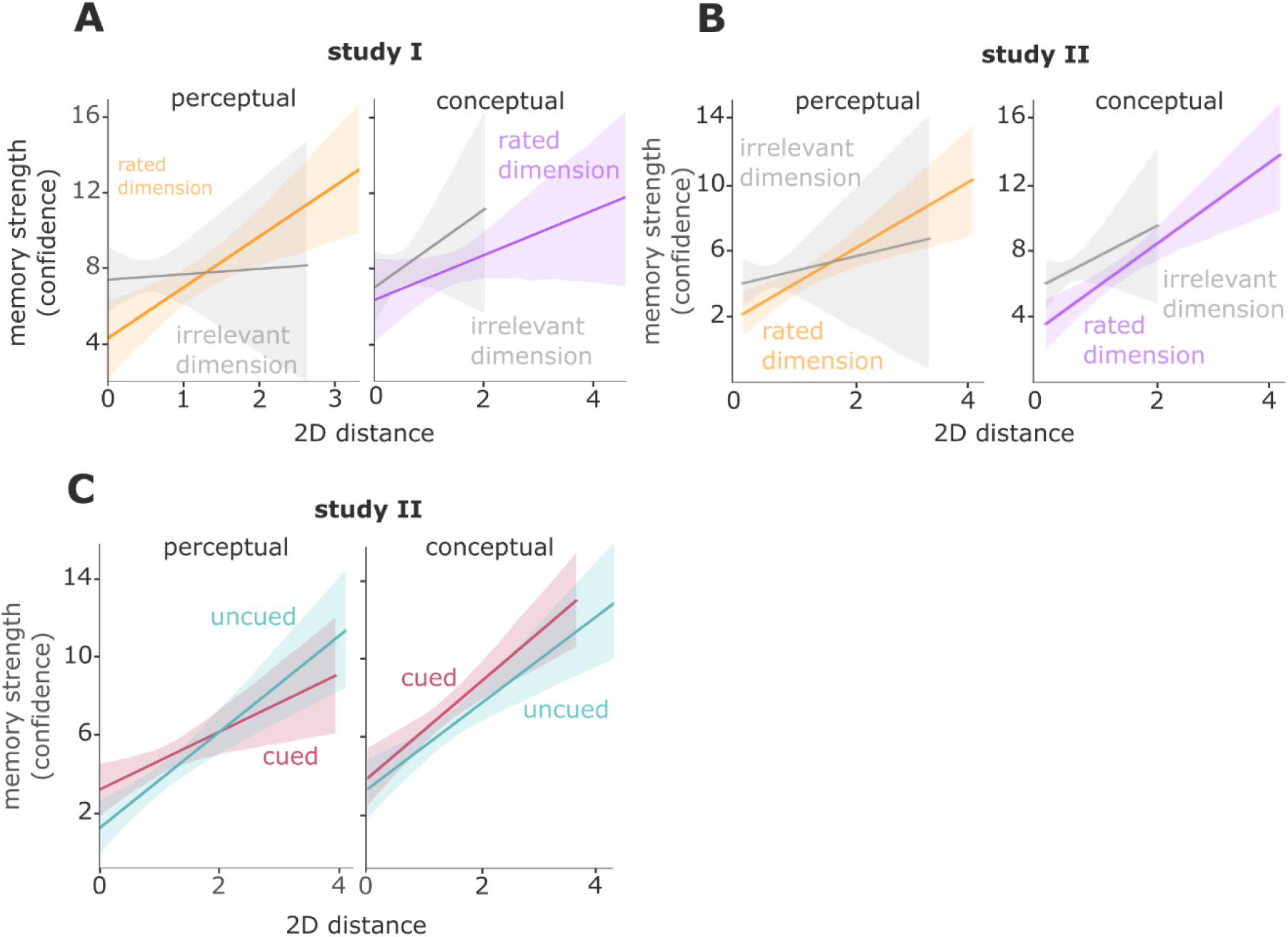
Representational distance affects subsequent memory regardless of perceptual/conceptual encoding space and cueing. (A) Study I: Encoding space did not affect the impact of representational distance on subsequent memory (interaction space x rated distance: z = 1.56, χ^2^_(2)_ = 2.89, p = 0.235). (B) Same for Study II (interaction space x rated distance: z = -0.63, χ^2^_(2)_ = 0.40, p = 0.820). (C) Study II: Cueing did not affect the impact of representational distance neither for perceptual (interaction cueing x rated distance [perceptual trials only]: z = 1.14, χ^2^_(2)_ = 1.30, p = 0.52) nor for conceptual encoding spaces (interaction cueing x rated distance [conceptual trials only]: z = - 0.11, χ^2^_(2)_ = 0.01, p = 0.99). Data are visualized after removing participant-wise estimated random effects. 95% confidence intervals.

**Supplementary Fig. 4.**
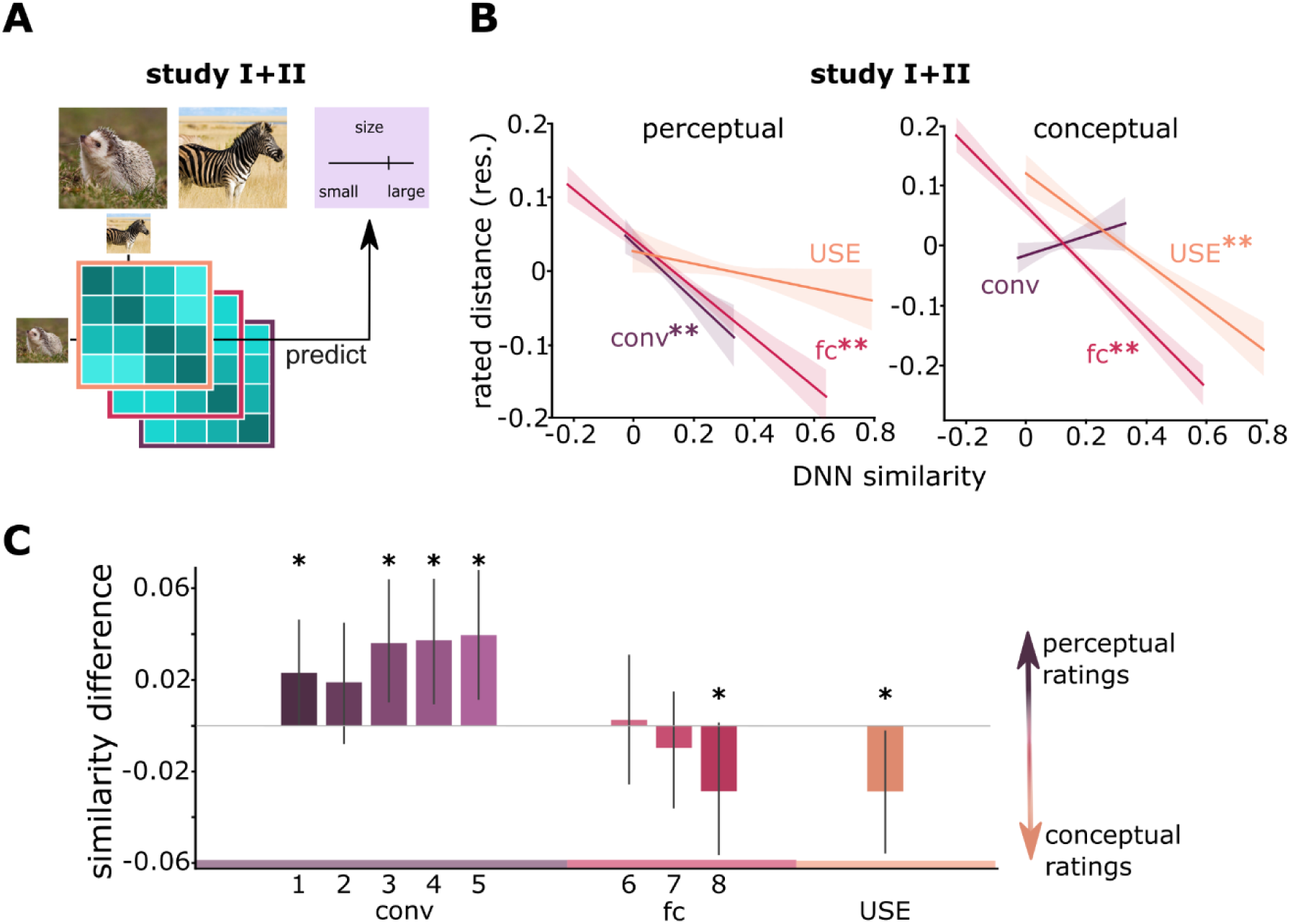
Layer-wise and averaged representational distances in Deep Neural Networks reflect behavioral similarity judgements. (A) We tested whether similarity judgements in the respective task-relevant perceptual and conceptual spaces corresponded to representational distances in specific DNN layers. To this end, we computed the similarity of each image pair in each network layer of a convolutional DNN and in the universal sentence encoder (USE) and used these similarities as predictors in a linear mixed model on rated distances of Studies I and II. (B) Using the layer averages (see Methods for more details) of convolutional (conv) and fully connected (fc) layers as well as the USE model similarities, we found that low-level perceptual and higher-order perceptual similarities predict rated distances during perceptual encoding. Reversely, conceptual ratings can be predicted by higher-order perceptual and semantic similarities (see Results for statistics). (C) In addition, we analyzed layer-wise similarities using paired t-tests (one-sided), FDR corrected for multiple comparisons. Effect sizes are measured by Cohen’s d. Indeed, representational distances in cDNN layers conv1 (t_101_ = 1.86, p = 0.049, d = 0.18, perceptual: M = 0.024, SD = 0.1, conceptual: M = 0.001, SD = 0.08), conv3 (t_101_ = 2.58, p = 0.017, d = 0.25, perceptual: M = 0.074, SD = 0.1, conceptual: M = 0.038, SD = 0.09), conv4 (t_101_ = 2.69, p = 0.017, d = 0.26, perceptual: M = 0.07, SD = 0.09, conceptual: M = 0.03, SD = 0.08) and conv5 (t_101_ = 2.88, p = 0.017, d = 0.28, perceptual: M = 0.08, SD = 09, conceptual: M = 0.04, SD = 0.08) predicted perceptual ratings better than conceptual ratings, while the reverse was true for fully-connected layer 8 (t_101_ = -1.93, p = 0.049, d = 0.19, perceptual: M = 0.10, SD = 0.09, conceptual: M = 0.13, SD = 0.10) and USE distances (t_101_ = -2.15, p = 0.037, d = 0.21, perceptual: M = 0.04, SD = 0.10, conceptual: M = 0.07, SD = 0.09) reflects the mean over all Spearman rho correlations of each DNN layer across all participant ratings). Data are visualized after removing participant-wise estimated random effects. 95% confidence intervals, error bars indicate SEM. *, p<0.05, **, p<0.01

**Supplementary Fig. 5.**
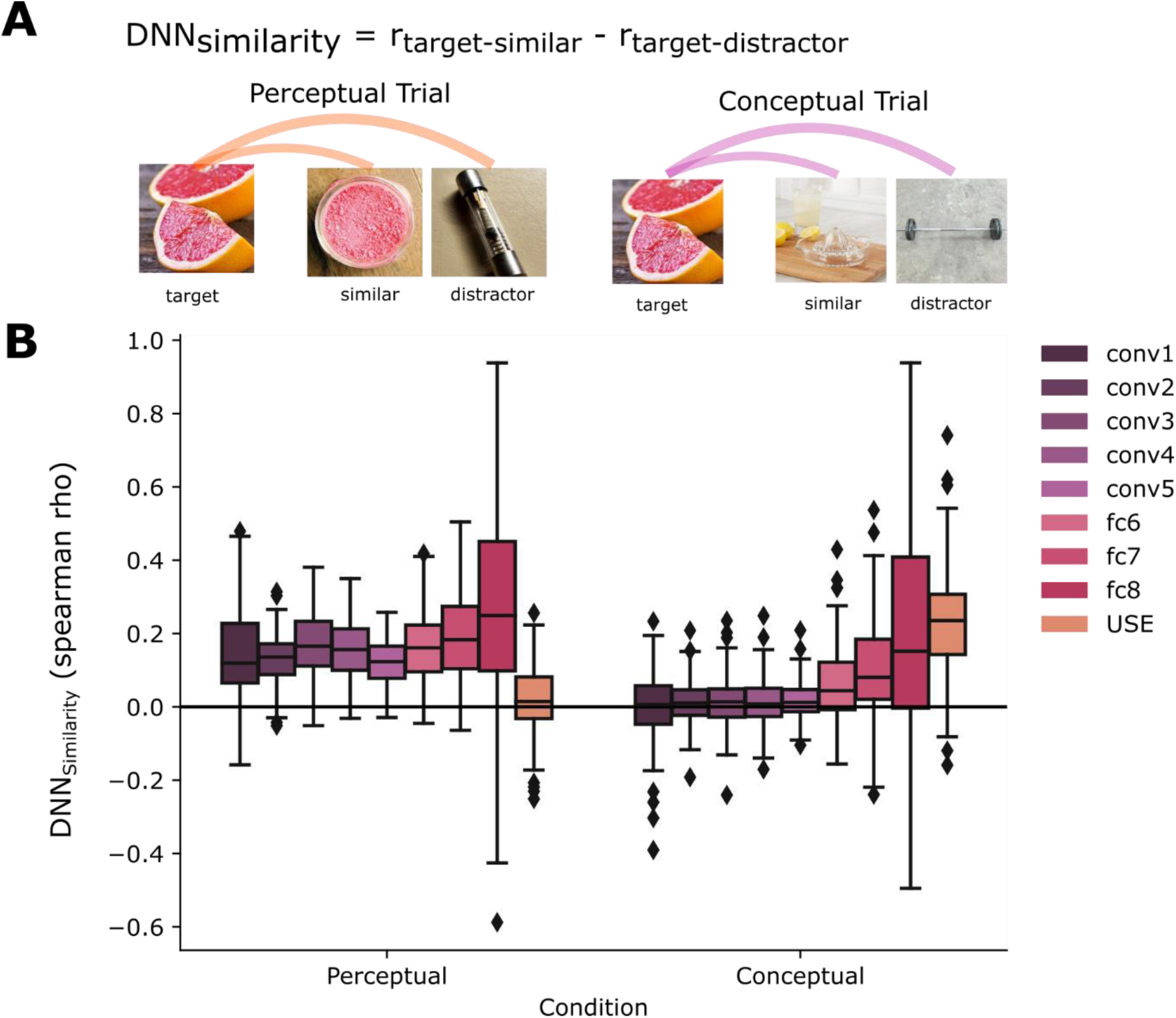
Layer-wise correspondence of representational distances in DNNs to binary ratings in Study III. (A) Exemplary target image with perceptual and conceptual choice options (similar choice, distractor choice). Lines indicate pairwise image similarities used to compute a triplet similarity score (d_similarity_). (B) Distribution of d_similarity_ scores across cDNN layers for conceptual and perceptual trials. Values larger than zero indicate higher DNN-based representational similarities of targets with the similar image as compared to the distractor image. For each layer and condition, boxplots depict the distribution of d_similarity_ scores across triplets. For perceptual triplets, d_similarity_ scores are significantly larger than zero for all cDNN layers, but not for the USE. For conceptual triplets, d_similarity_ scores are significantly larger than zero from conv2 until and including the USE model. Center line indicates median, box limits indicate upper and lower quartiles, whiskers indicate 1.5 interquartile range, points represent outliers.

**Supplementary Fig. 6.**
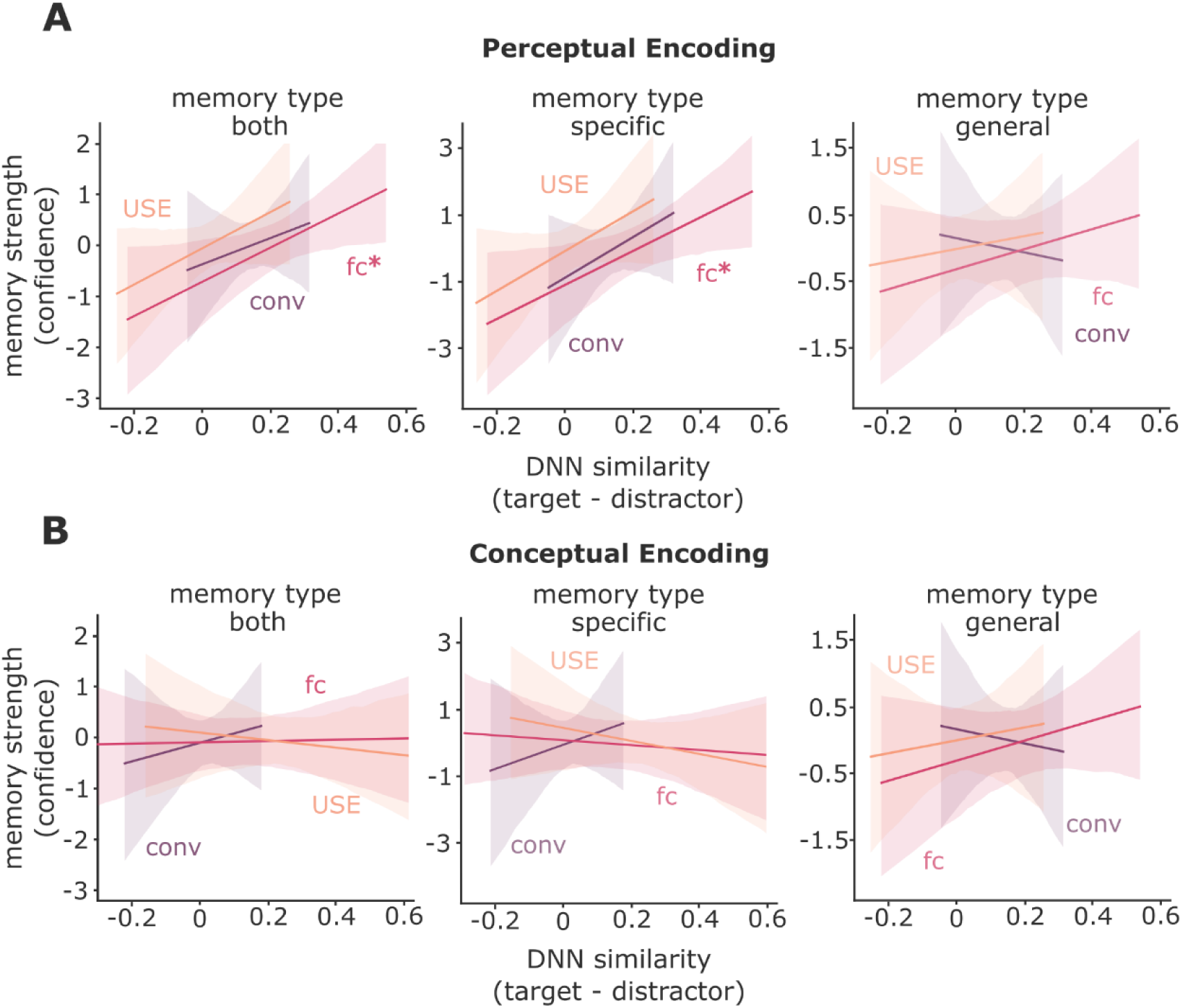
Effects of DNN-based representational similarities on memory strength during the general and the specific memory task. (A) Prediction of memory strength by DNN similarities during perceptual choices. Left: Collapsed across memory types, representational distances in fully-connected layers but not convolutional layers (conv) and not semantic distances (USE-based) predict memory strength (see Results section). Middle: Memory strength in the specific retrieval task is predicted by fully-connected (: z = 2.258, χ^2^_(1)_ = 5.09, p = 0.024) but not convolutional or semantic (conv: z = 1.138, χ^2^_(1)_ = 1.29, p = 0.255; USE: z = 1.567, χ^2^_(1)_ = 2.45, p = 0.117; likelihood ratio test) DNN similarities during perceptual encoding. Right: Memory in the general memory test is independent of DNN similarities during perceptual encoding (conv: z = -0.293, χ^2^_(1)_ = 0.09, p = 0.769; fc: z = 0.977, χ^2^_(1)_ = 0.95, p = 0.328; USE: z = 0.373, χ^2^_(1)_ = 0.14, p = 0.709). (B) Prediction of memory strength by DNN similarities during conceptual encoding. Left: Collapsed across memory types, DNN-predicted similarities do not predict subsequent memory (see Results section). Middle: Memory strength during specific retrieval cannot be predicted by DNN similarities (conv: z = 0.667, χ^2^_(1)_ = 0.45, p = 0.504; fc: z = -0.404, χ^2^_(1)_ = 0.16, p = 0.686; USE: z = -0.807, χ^2^_(1)_ = 0.65, p = 0.419; likelihood ratio test). Right: Memory strength during general retrieval cannot be predicted by DNN similarities. For studies I and II, where task-demands did not differ between general and specific retrieval, we found no effect of DNN similarities on memory strengths (Study I+II: conv: z = 0.49, χ^2^_(1)_ = 1.34, p = 0.247; fc: z = 0.94, χ^2^_(1)_ =1.00, p = 0.317; USE: z = 0.87, χ^2^_(1)_ = 1.90, p = 0.167; likelihood ratio test). Data are visualized after removing participant-wise estimated random effects. 95% confidence intervals. *, p<0.05

**Supplementary Fig. 7.**
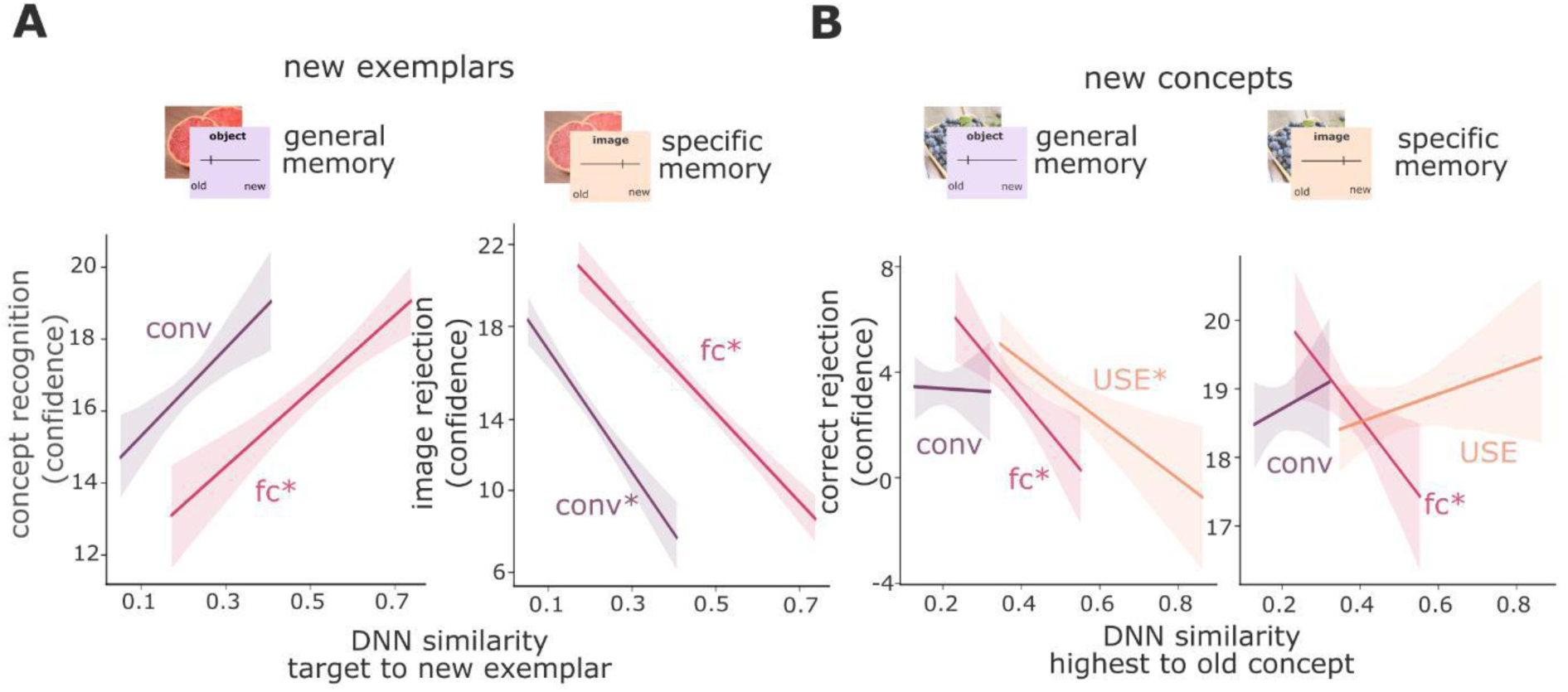
Different formats contribute to memory strength and correct rejection performance depending on task instructions at retrieval. (A) We computed the similarity of each new exemplar image to the old image of the same category and extracted the similarity value as the DNN similarity (for each DNN layer). Using a linear mixed model including both the main effects of DNN layers and task, we found significant effects of retrieval instruction with distances in low-level perceptual space (z = -2.22, χ^2^_(1)_ = 13.30, p = 0.0002; likelihood ratio test) and higher-order perceptual space (z = -7.89, χ^2^_(1)_ = 70.34, p<0.0001; likelihood ratio test) indicating that representational distances between new and previously presented items exerted different effects depending on retrieval tasks. Correct rejection performance for new exemplars in the specific retrieval task depended on similarities of both low-level perceptual (z =-2.51, χ^2^_(1)_ = 6.34, p = 0.011; likelihood ratio test) and higher-order perceptual (z = -7.16, χ^2^_(1)_ = 50.97, p<0.0001; likelihood ratio test) formats, while memory strength (confidence) for old concepts during general retrieval depended on similarity of higher-order (z = 4.40, χ^2^_(1)_ = 19.37, p<0.0001; likelihood ratio test) but not low-level perceptual formats (z = 0.51, χ^2^_(1)_ =0.26, p = 0.6090; likelihood ratio test). (Note that semantic formats could not be considered in this analysis as new exemplars always belonged to the same concept.) (B) For new concepts we calculated the semantic similarity to all encoding concepts and extracted the highest value (highest similarity of new concept – all old images/concepts): Using a linear mixed model including DNN similarities as predictors, we found that higher-order perceptual (z =-2.78, χ^2^_(1)_ = 7.73, p = 0.005; likelihood ratio test) but not low-level perceptual (z =1.39, χ^2^_(1)_ =1.95, p = 0.162; likelihood ratio test) or USE (z =0.50, χ^2^_(1)_ = 0.25, p = 0.615; likelihood ratio test) similarities predicted performance during specific retrieval. During general retrieval, both higher-order perceptual (z =-4.31, χ^2^_(1)_ = 18.57, p < 0.0001; likelihood ratio test) and semantic similarities (z =-3.96, χ^2^_(1)_ = 15.65, p < 0.0001; likelihood ratio test) but not low-level perceptual similarities (z =1.14, χ^2^_(1)_ = 1.32, p = 0.250; likelihood ratio test) predicted rejection of new concepts. Data are visualized after removing participant-wise estimated random effects. 95% confidence intervals. *, p<0.05

**Supplementary Fig. 8.**
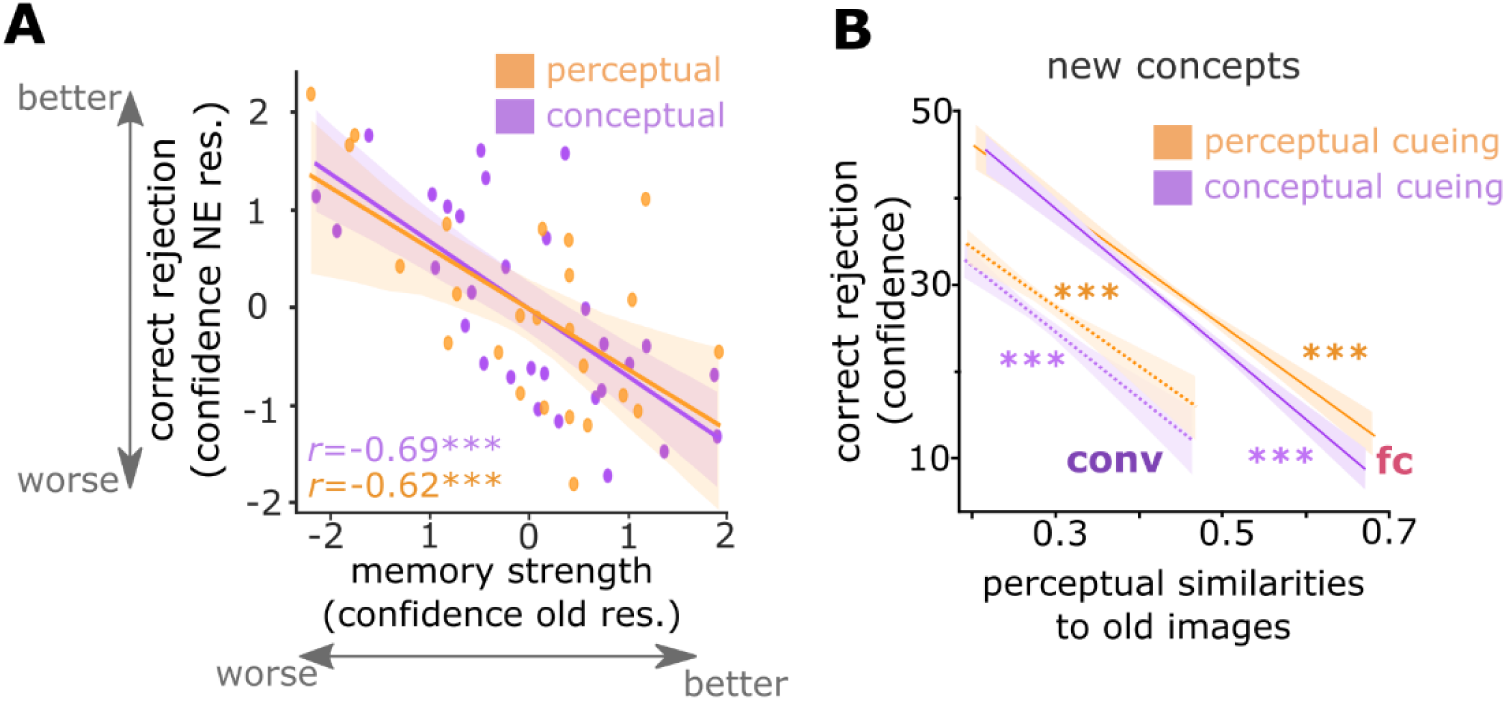
Conceptual cueing specifically influences semantic memory. (A) We tested whether participants who showed higher recognition memory performance also showed a decrease in rejection performance of new exemplars because they generally used a more liberal criterion during memory decisions. To this end, we calculated Pearson’s correlations on participants’ mixed model residuals. We found that participants in both conceptual (*r* = -0.69, p<0.001) and perceptual (*r* = -0.62, p<0.001) cueing conditions show this effect, i.e., those who performed better in identifying old images made more errors for new exemplars. (B) Since conceptual cueing impaired the rejection of new concepts that were semantically highly similar to old concepts, we tested whether cueing affected the rejection of perceptually similar new concepts as well. To this end – analogously to our analysis on semantic similarities (Fig. 5d) – we computed two linear mixed models (one for perceptual cueing and one for conceptual cueing) using perceptual similarities from the cDNN as predictors (highest similarity new concept – old images). Perceptual features predicted correct rejection confidence in both perceptual (conv: z = -6.64, χ^2^_(1)_ = 43.84, p < 0.001; fc: z = -13.71, χ^2^_(1)_ = 183.88, p < 0.001) and conceptual cueing conditions (conv: z = -7.21, χ^2^_(1)_ = 51.69, p < 0.001; fc: z = -15.46, χ^2^_(1)_ = 233.53, p < 0.001). Thus, low-level perceptual (conv) and higher-order perceptual (fc) similarities impaired rejection performance following both perceptual and conceptual cueing. Data are visualized after removing participant-wise estimated random effects. 95% confidence intervals. ***, p<0.001

**Supplementary Table 1.**
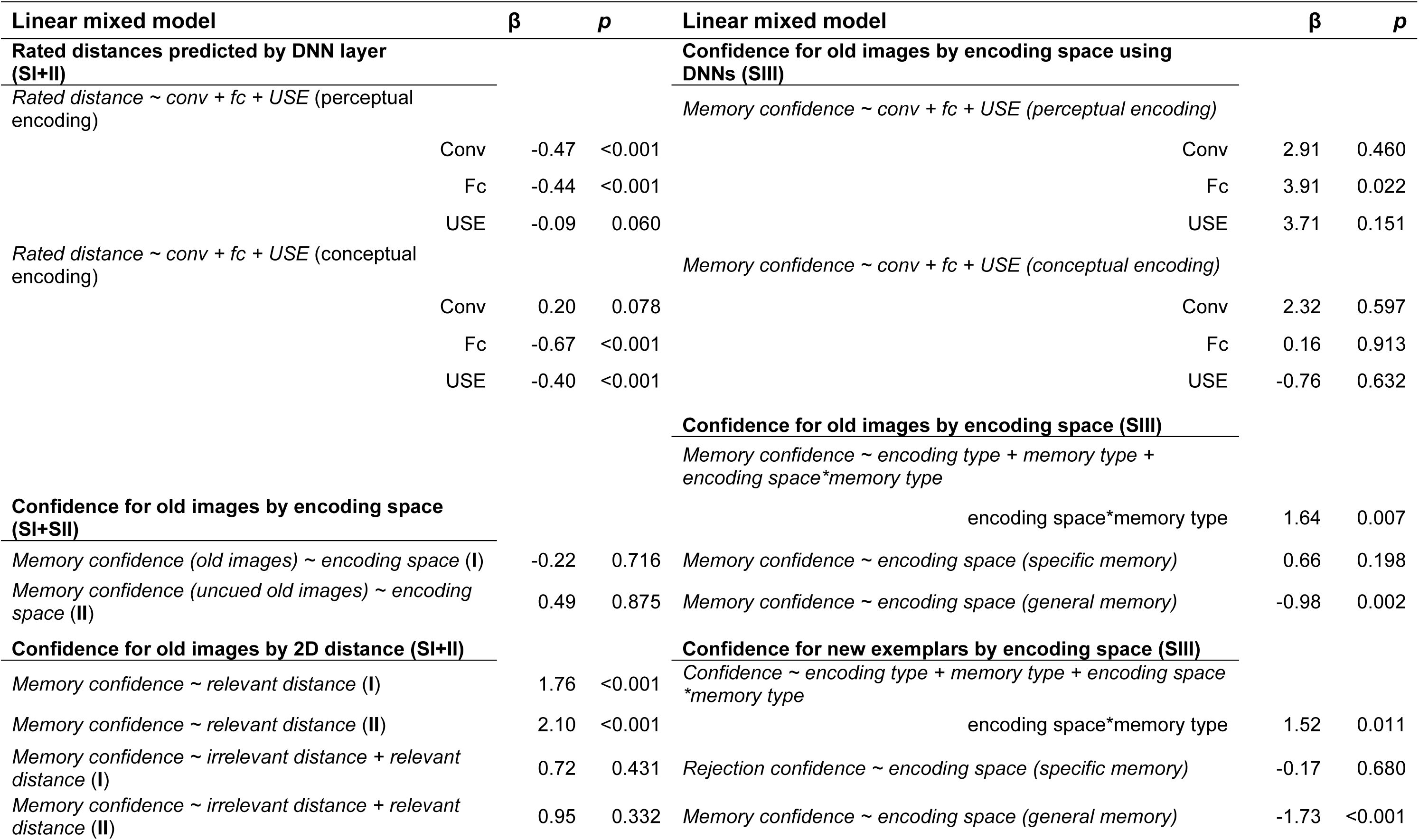
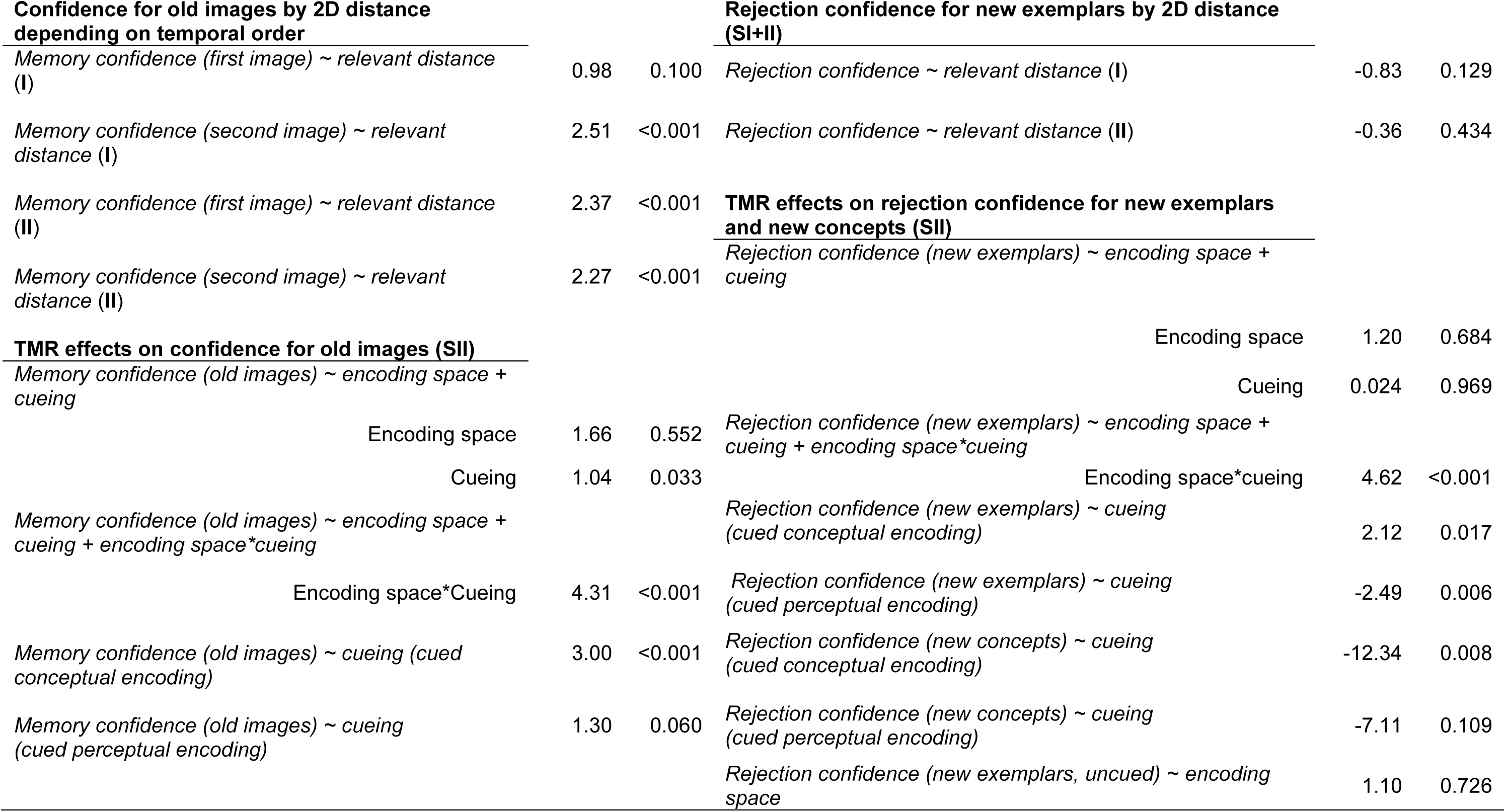
Linear mixed models and their beta weights from main results. Corresponding Beta weights to the z values reported in the main results. All models include a random participant intercept (1|participant).

| Linear mixed model | $\beta$ | $p$ | Linear mixed model | $\beta$ | $p$ |
| --- | --- | --- | --- | --- | --- |
| **Rated distances predicted by DNN layer (SI+II)** |  |  | **Confidence for old images by encoding space using DNNs (SIII)** |  |  |
| *Rated distance ~ conv + fc + USE (perceptual encoding)* |  |  | *Memory confidence ~ conv + fc + USE (perceptual encoding)* |  |  |
| Conv | -0.47 | <0.001 | Conv | 2.91 | 0.460 |
| Fc | -0.44 | <0.001 | Fc | 3.91 | 0.022 |
| USE | -0.09 | 0.060 | USE | 3.71 | 0.151 |
| *Rated distance ~ conv + fc + USE (conceptual encoding)* |  |  | *Memory confidence ~ conv + fc + USE (conceptual encoding)* |  |  |
| Conv | 0.20 | 0.078 | Conv | 2.32 | 0.597 |
| Fc | -0.67 | <0.001 | Fc | 0.16 | 0.913 |
| USE | -0.40 | <0.001 | USE | -0.76 | 0.632 |
| **Confidence for old images by encoding space (SI+SII)** |  |  | **Confidence for old images by encoding space (SIII)** |  |  |
| *Memory confidence (old images) ~ encoding space (I)* |  |  | *Memory confidence ~ encoding type + memory type + encoding space\*memory type* |  |  |
|  | -0.22 | 0.716 | encoding space*memory type | 1.64 | 0.007 |
| *Memory confidence (uncued old images) ~ encoding space (II)* |  |  | *Memory confidence ~ encoding space (specific memory)* |  |  |
|  | 0.49 | 0.875 |  | 0.66 | 0.198 |
| **Confidence for old images by 2D distance (SI+II)** |  |  | *Memory confidence ~ encoding space (general memory)* |  |  |
| *Memory confidence ~ relevant distance (I)* |  |  |  | -0.98 | 0.002 |
|  | 1.76 | <0.001 | **Confidence for new exemplars by encoding space (SIII)** |  |  |
| *Memory confidence ~ relevant distance (II)* |  |  | *Confidence ~ encoding type + memory type + encoding space\*memory type* |  |  |
|  | 2.10 | <0.001 | encoding space*memory type | 1.52 | 0.011 |
| *Memory confidence ~ irrelevant distance + relevant distance (I)* |  |  | *Rejection confidence ~ encoding space (specific memory)* |  |  |
|  | 0.72 | 0.431 |  | -0.17 | 0.680 |
| *Memory confidence ~ irrelevant distance + relevant distance (II)* |  |  | *Memory confidence ~ encoding space (general memory)* |  |  |
|  | 0.95 | 0.332 |  | -1.73 | <0.001 |

Supp Table 1. continued
| Confidence for old images by 2D distance<br>depending on temporal order |  |  | Rejection confidence for new exemplars by 2D distance<br>(SI+II) |  |  |
| --- | --- | --- | --- | --- | --- |
| Memory confidence (first image) ~ relevant distance (I) | 0.98 | 0.100 | Rejection confidence ~ relevant distance (I) | -0.83 | 0.129 |
| Memory confidence (second image) ~ relevant distance (I) | 2.51 | <0.001 | Rejection confidence ~ relevant distance (II) | -0.36 | 0.434 |
| Memory confidence (first image) ~ relevant distance (II) | 2.37 | <0.001 | **TMR effects on rejection confidence for new exemplars and new concepts (SII)** |  |  |
| Memory confidence (second image) ~ relevant distance (II) | 2.27 | <0.001 | Rejection confidence (new exemplars) ~ encoding space + cueing |  |  |
| **TMR effects on confidence for old images (SII)** |  |  | Encoding space | 1.20 | 0.684 |
| Memory confidence (old images) ~ encoding space + cueing |  |  | Cueing | 0.024 | 0.969 |
| Encoding space | 1.66 | 0.552 | Rejection confidence (new exemplars) ~ encoding space + cueing + encoding space*cueing |  |  |
| Cueing | 1.04 | 0.033 | Encoding space*cueing | 4.62 | <0.001 |
| Memory confidence (old images) ~ encoding space + cueing + encoding space*cueing |  |  | Rejection confidence (new exemplars) ~ cueing (cued conceptual encoding) | 2.12 | 0.017 |
| Encoding space*Cueing | 4.31 | <0.001 | Rejection confidence (new exemplars) ~ cueing (cued perceptual encoding) | -2.49 | 0.006 |
| Memory confidence (old images) ~ cueing (cued conceptual encoding) | 3.00 | <0.001 | Rejection confidence (new concepts) ~ cueing (cued conceptual encoding) | -12.34 | 0.008 |
| Memory confidence (old images) ~ cueing (cued perceptual encoding) | 1.30 | 0.060 | Rejection confidence (new concepts) ~ cueing (cued perceptual encoding) | -7.11 | 0.109 |
|  |  |  | Rejection confidence (new exemplars, uncued) ~ encoding space | 1.10 | 0.726 |

**Supplementary Table 2.**
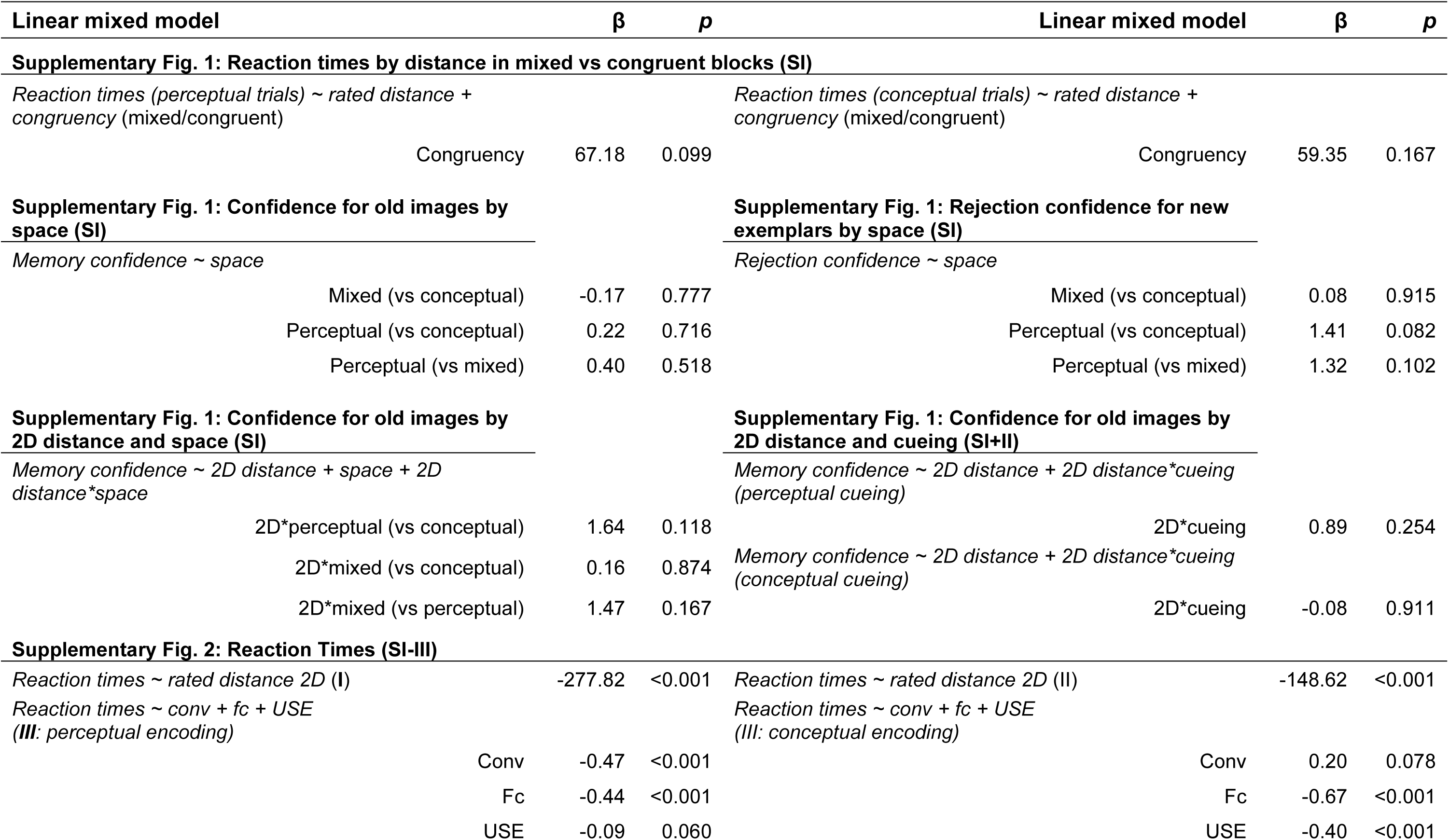
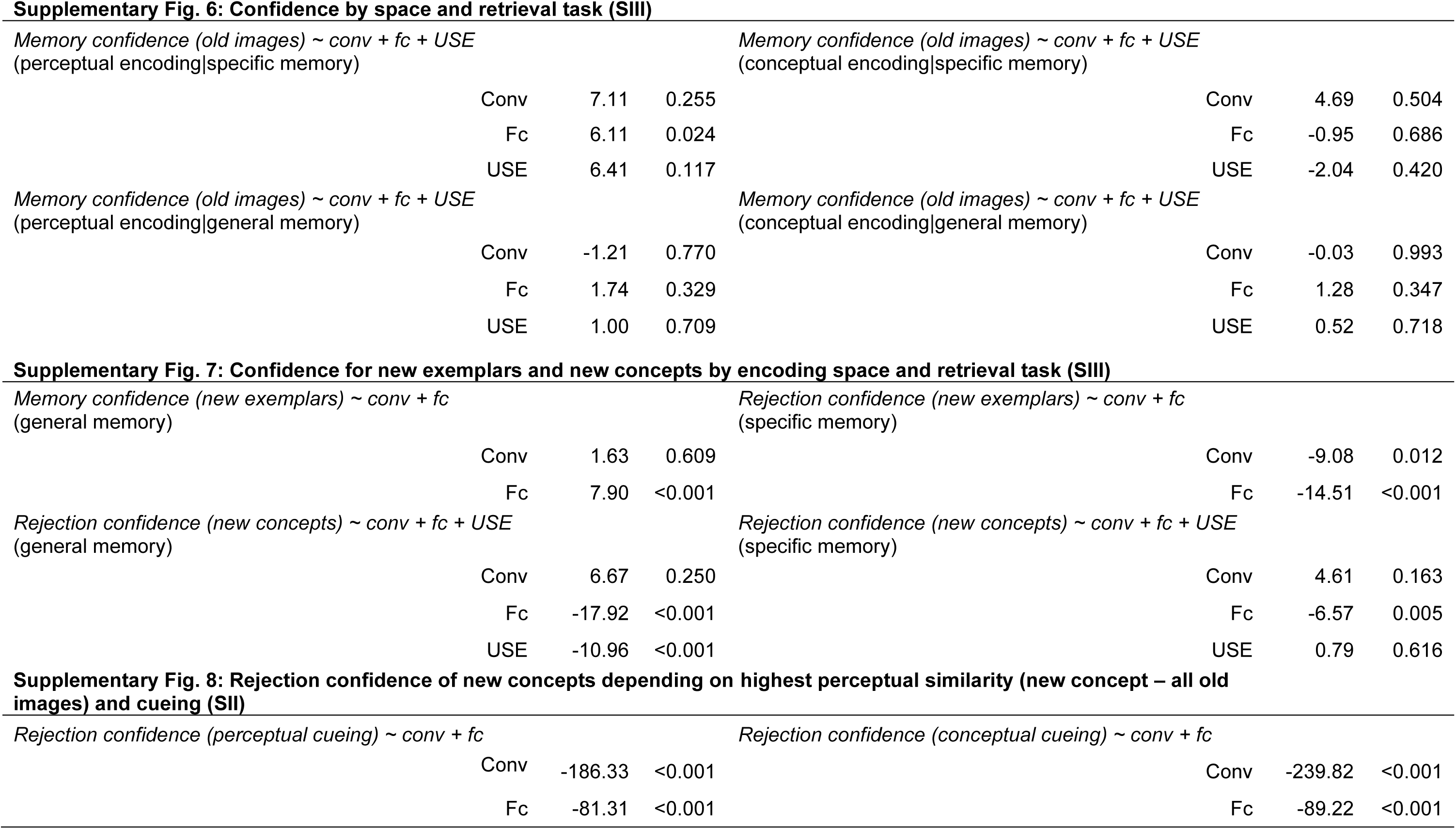
Linear mixed models and their beta weights from Supplementary Analyses. Corresponding Beta weights to the z values reported in the Supplementary Analyses. All models include a random participant intercept (1|participant).

| Linear mixed model | $\beta$ | $p$ | Linear mixed model | $\beta$ | $p$ |
| --- | --- | --- | --- | --- | --- |
| **Supplementary Fig. 1: Reaction times by distance in mixed vs congruent blocks (SI)** |  |  |  |  |  |
| *Reaction times (perceptual trials) ~ rated distance + congruency (mixed/congruent)* |  |  | *Reaction times (conceptual trials) ~ rated distance + congruency (mixed/congruent)* |  |  |
| Congruency | 67.18 | 0.099 | Congruency | 59.35 | 0.167 |
| **Supplementary Fig. 1: Confidence for old images by space (SI)** |  |  | **Supplementary Fig. 1: Rejection confidence for new exemplars by space (SI)** |  |  |
| *Memory confidence ~ space* |  |  | *Rejection confidence ~ space* |  |  |
| Mixed (vs conceptual) | -0.17 | 0.777 | Mixed (vs conceptual) | 0.08 | 0.915 |
| Perceptual (vs conceptual) | 0.22 | 0.716 | Perceptual (vs conceptual) | 1.41 | 0.082 |
| Perceptual (vs mixed) | 0.40 | 0.518 | Perceptual (vs mixed) | 1.32 | 0.102 |
| **Supplementary Fig. 1: Confidence for old images by 2D distance and space (SI)** |  |  | **Supplementary Fig. 1: Confidence for old images by 2D distance and cueing (SI+II)** |  |  |
| *Memory confidence ~ 2D distance + space + 2D distance\*space* |  |  | *Memory confidence ~ 2D distance + 2D distance\*cueing (perceptual cueing)* |  |  |
| 2D*perceptual (vs conceptual) | 1.64 | 0.118 | 2D*cueing | 0.89 | 0.254 |
| 2D*mixed (vs conceptual) | 0.16 | 0.874 | *Memory confidence ~ 2D distance + 2D distance\*cueing (conceptual cueing)* |  |  |
| 2D*mixed (vs perceptual) | 1.47 | 0.167 | 2D*cueing | -0.08 | 0.911 |
| **Supplementary Fig. 2: Reaction Times (SI-III)** |  |  |  |  |  |
| *Reaction times ~ rated distance 2D (I)* | -277.82 | <0.001 | *Reaction times ~ rated distance 2D (II)* | -148.62 | <0.001 |
| *Reaction times ~ conv + fc + USE (III: perceptual encoding)* |  |  | *Reaction times ~ conv + fc + USE (III: conceptual encoding)* |  |  |
| Conv | -0.47 | <0.001 | Conv | 0.20 | 0.078 |
| Fc | -0.44 | <0.001 | Fc | -0.67 | <0.001 |
| USE | -0.09 | 0.060 | USE | -0.40 | <0.001 |

|  |  |  |
| --- | --- | --- |
| Conv | 7.11 | 0.255 |
| Fc | 6.11 | 0.024 |
| USE | 6.41 | 0.117 |

|  |  |  |
| --- | --- | --- |
| Conv | -1.21 | 0.770 |
| Fc | 1.74 | 0.329 |
| USE | 1.00 | 0.709 |

|  |  |  |
| --- | --- | --- |
| Conv | 4.69 | 0.504 |
| Fc | -0.95 | 0.686 |
| USE | -2.04 | 0.420 |

|  |  |  |
| --- | --- | --- |
| Conv | -0.03 | 0.993 |
| Fc | 1.28 | 0.347 |
| USE | 0.52 | 0.718 |

|  |  |  |
| --- | --- | --- |
| Conv | 1.63 | 0.609 |
| Fc | 7.90 | <0.001 |

|  |  |  |
| --- | --- | --- |
| Conv | 6.67 | 0.250 |
| Fc | -17.92 | <0.001 |
| USE | -10.96 | <0.001 |

|  |  |  |
| --- | --- | --- |
| Conv | -9.08 | 0.012 |
| Fc | -14.51 | <0.001 |

|  |  |  |
| --- | --- | --- |
| Conv | 4.61 | 0.163 |
| Fc | -6.57 | 0.005 |
| USE | 0.79 | 0.616 |

|  |  |  |
| --- | --- | --- |
| Conv | -186.33 | <0.001 |
| Fc | -81.31 | <0.001 |

*Rejection confidence (conceptual cueing) ~ conv + fc*
|  |  |  |
| --- | --- | --- |
| Conv | -239.82 | <0.001 |
| Fc | -89.22 | <0.001 |

